# Horizontally Tiled Network of Cortico-Basal Ganglia Modules Performs Reinforcement Learning

**DOI:** 10.1101/2025.06.18.660381

**Authors:** Kojiro Hirokane, Ryo Takehara, Tetsuji Kita, Shunya Onoi, Soma Matsushita, Takashi Kitsukawa

## Abstract

The neocortex and basal ganglia nuclei are connected along regions that share the same topography and are arranged side-by-side. Inspired by the anatomical characteristics of the cerebrum, we developed a network in which the modules of the neocortex-basal ganglia unit were arranged in a horizontally tiled manner. By applying this network to reinforcement learning tasks, we demonstrated that reinforcement learning can be achieved through horizontal signals passing between modules. Each module not only performs its calculation but also provides signals to adjacent modules. This lateral transmission takes advantage of the differences in the projection ranges of the three basal ganglia pathways, the direct, indirect, and hyperdirect pathways, which have been examined in physiologic studies. We found that these differences enabled temporal-difference-error computations. This study proposes a novel strategy for information processing based on neocortical-basal ganglia circuits, highlighting the computational significance of their anatomically and physiologically clarified features.

## Introduction

Reinforcement learning (RL) is a fundamental mechanism through which we learn from our environment by taking actions and receiving feedback in the form of rewards or punishments. This mechanism helps guide future behavior by selecting the most appropriate actions based on past experiences. This process is essential for decision making, behavioral adaptation, and habit formation in daily life.

The basal ganglia play a central role in RL by processing rewards and punishments. The striatum, a component of the basal ganglia, is involved in the representation of the expected future rewards associated with specific actions, known as ‘Q-value’^1^. This allows us to evaluate and select actions that maximize rewards over time. Striatal neurons receive input from dopaminergic neurons in the substantia nigra pars compacta carrying essential information related to RL, such as reward signals and reward prediction errors^2,3^. Dopamine plays a critical role in updating Q-values by signaling when an outcome is better or worse than expected, thus reinforcing behaviors that lead to rewards. Despite strong evidence for the involvement of the basal ganglia in RL, the neural circuits responsible for executing RL processes and the exact pathways and interactions between the nuclei involved remain unclear.

Key anatomical and physiological features of the basal ganglia have been delineated. The cortico-basal ganglia network is formed by three distinct pathways connecting the neocortex and substantia nigra pars reticulata (SNr). The hyperdirect pathway exerts an excitatory influence on the SNr via the subthalamic nuclei (STN). The direct pathway exerts an inhibitory influence on the SNr through the striatal direct pathway neurons. The indirect pathway exerts an excitatory influence on the SNr through striatal indirect pathway neurons and the external segment of the globus pallidus (GPe). These nuclei all have topographic maps corresponding to body parts^4,5^, which suggests that they are interconnected in accordance with the maps.

In addition, physiological studies have demonstrated that these three pathways differ in their projection ranges in the SNr^6–9^. By recording the range of responses in the SNr to stimuli applied to the cortex, Nambu et al.^7^ and Ozaki et al.^9^ demonstrated that the direct pathway projects locally in the SNr/GPi, whereas the hyperdirect and indirect pathways project across a wider range in the SNr/GPi. These differences in the projection ranges appear to be partially at odds with the findings of the topographic maps for all nuclei.

Herein, we propose a novel view of the basal ganglia, suggesting that the cortico-basal ganglia organization consist of horizontally tiled modular microcircuits (BG-modules) of topographically matched neurons throughout the nuclei. We demonstrate that this topographical architecture helps the network solve RL tasks, using specific inter-module connections inferred from the differences in the projection ranges of three distinct pathways to represent the value in RL tasks. The results of this study should advance our understanding of the pathophysiology of diseases characterized by functional abnormalities of the basal ganglia.

## Results

### Tiled network is a neural network inspired by the basal ganglia circuit

In this study, we focused on whether the modular structure of a tiled network helps RL by evaluating its function using a maze task, a type of RL task. RL is a paradigm of machine learning where an agent interacts with the environment and learns to select optimal actions based on feedback resulting from its actions (see Sutton & Barto, 2018)^10^ for a detailed treatment of RL. Note that our notation differs slightly from theirs. The agent must be a system that receives observations from the environment (e.g., its current position) and outputs actions. In other words, if we denote the agent’s state at time T = *t* as *s*_*t*_ ∈ ***S*** and the action chosen by the agent as *a*_*t*_ ∈ ***A***, the state *s*_*t*+1_ at the next time T = *t+1* is determined by the state *s*_*t*_ and action *a*_*t*_ at time T = *t* as follows:

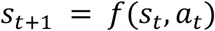

Here, ***S*** refers to the set of all possible states in the environment and *A* refers to the set of all possible actions that the agent can take. By determining the actions and transitioning states, as described above, the agent explores the environment and maximizes the rewards. Specifically, we implemented an agent using a tiled network as follows.

We first hypothesized a parallel cortico-basal ganglia circuit based on the known anatomical and physiological connections between the major nuclei in the basal ganglia (Fig. 1A). The cortico-basal ganglia circuit is composed of the cortex, striatum, and external and internal segments of the globus pallidus (GPe and GPi), SNr, STN, and substantia nigra compacta (SNc). Within the striatum, D1 receptor-expressing striatal neurons (striatal D1 SPNs) project directly to the SNr/GPi, and D2 receptor-expressing striatal neurons (striatal D2 SPNs) project to the GPe and then to the SNr/GPi, making this thus an indirect projection system. We use human anatomical terminology here; although the basal ganglia are highly conserved across species^11^, the anatomical terms may differ. We modeled each nucleus as a single firing-rate neuron, representing the average activity level of the nucleus.

**Figure 1.**
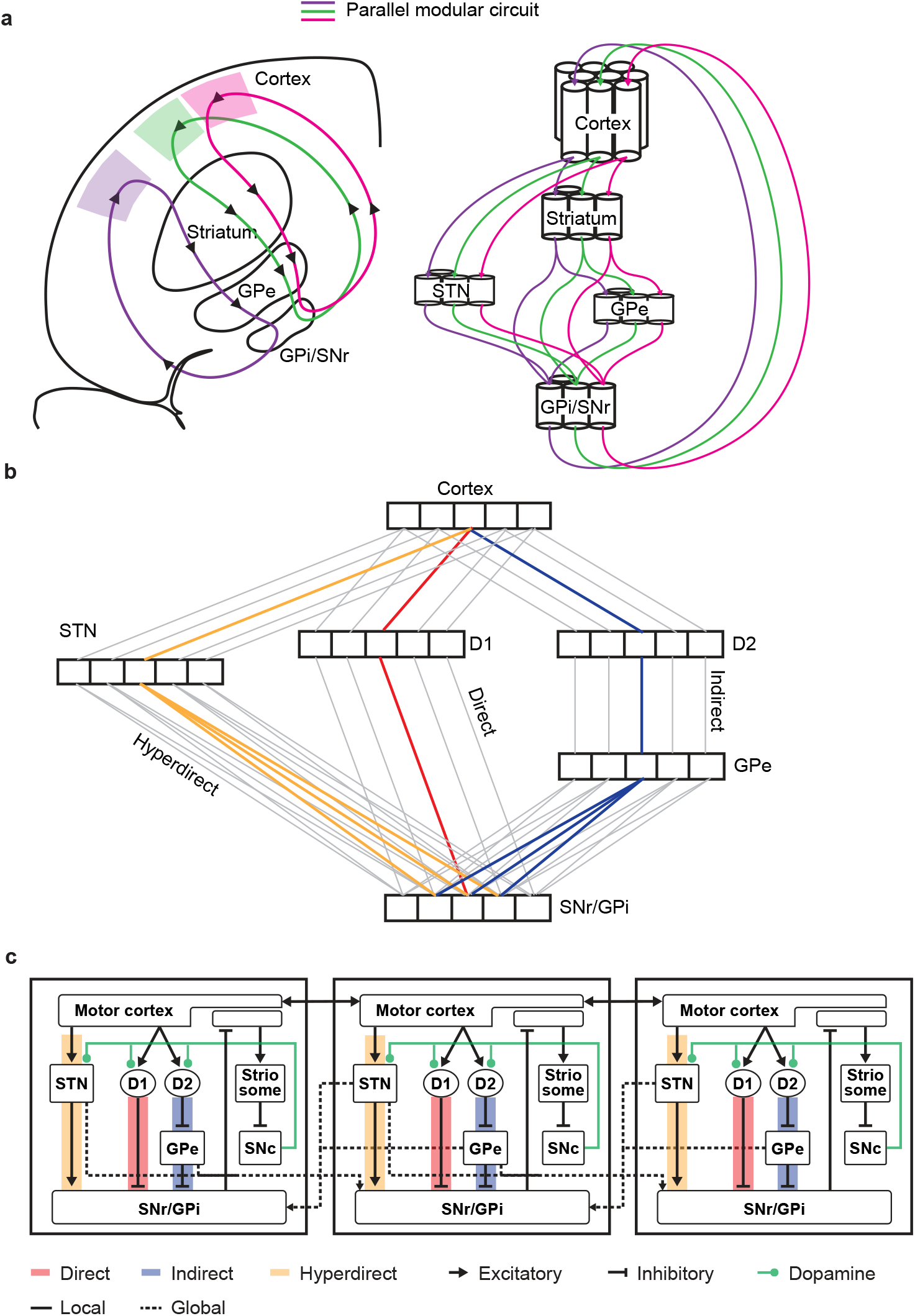
Implementation of a tiled network composed of tiled BG-modules. A. Abstract of the neural circuit assumed in this study: the cortico-basal ganglia circuit forms a parallel modular circuit. B. Direct pathway (Cortex→striatal D1 SPNs→SNr/GPi) within the basal ganglia have a narrower projection range, whereas hyperdirect pathway (Cortex→STN→SNr/GPi) and indirect pathway (Cortex→striatal D2 SPNs→SNr/GPi) have a wider range. C. Diagram of connections in horizontally tiled BG-modules for maze task (solid line: intra-module connections, dotted line: inter-module connections).

The cortico-basal ganglia circuit contains multiple pathways, each with either excitatory or inhibitory effects on the output nucleus of the basal ganglia (SNr/GPi). Consequently, they have opposing effects on the downstream feedback projections to the cortex via the thalamus. There are three main pathways in the basal ganglia circuit: the hyperdirect pathway (Cortex → STN → SNr/GPi), the direct pathway (cortex → striatal D1 SPNs→ SNr/GPi), and the indirect pathway (cortex → striatal D2 SPNs→ GPe → SNr/GPi)^12,13^. When considering their impact on the cortex, signals through the direct pathway are net excitatory, whereas those through the hyperdirect and indirect pathways are net inhibitory. We did not include the thalamus in our model because its projections are excitatory, meaning that the nucleus does not switch the excitatory-inhibitory effect within the circuit. Therefore, excluding the thalamus from our model did not compromise the essential computational properties of the circuit. Based on these insights, we implemented these three pathways by connecting the nuclei by weight for each parallel circuit.

We also implemented a dopamine input into the striatum. In addition to inputs from the cortex, the striatum receives projections from dopamine neurons in the SNc. When the striatum receives dopaminergic input, the activity of striatal D1 neurons increases, whereas that of striatal D2 neurons decreases^14,15^. Importantly, these opposing effects lead to increased cortical excitation when dopamine levels in the striatum increase, provided that cortical inputs remain constant. Changes in activity also strengthen synapses between cortical and D1 neurons, while weakening connections between cortical and D2 neurons^16^. This dynamic modulation enables the basal ganglia to adjust to the cortical influences of excitation and inhibition in response to dopamine input. This supports our model’s use of two-factor Hebbian plasticity rules based on presynaptic glutamatergic inputs and dopaminergic innervation^16,17^.

Moreover, in addition to the matrix compartment, the striatum contains a compartment called the striosome. Each compartment has distinct functions^18,19^. For instance, the striosome directly regulates dopamine release^20,21^. Therefore, in addition to the matrix compartment, we implemented a striosomal compartment and connected it directly to the SNc.

Furthermore, as the most critical part of our model, we hypothesized that hyperdirect and indirect pathways have wider projection ranges, extending their projections to neighboring parallel circuits. In contrast, the direct pathway had a narrower projection range and was connected only within a parallel circuit (Fig. 1B). This was based on a physiological study in which responses to cortical stimulation via the hyperdirect and indirect pathways were recorded from a wider area in the SNr/GPi, whereas those via the direct pathway were recorded only from a narrower area in the SNr/GPi^9^. Given these facts, we implemented a horizontally tiled cortico-basal ganglia module (BG-module), with each module containing the aforementioned nuclei that formed three pathways (Fig. 1C). Specifically, the direct pathway (striatal D1 SPNs → SNr/GPi) was confined to intra-module projections, whereas the hyperdirect pathway (STN → SNr/GPi) and indirect pathway (GPe → SNr/GPi) were both intra- and inter-module projections to adjacent modules (Fig. 1C).

### Learning algorithm of the tiled network in a 1D maze task

We first used a 1D maze task to evaluate the core feature of our model that the agent can learn by gradually changing the activity levels of the subcortical nuclei within each module, driven by inter-module interactions via the horizontally connected network. In the maze task, the agent receives its current position from the environment and selects an action to move left or right (Fig. 2A). We configured the BG-modules corresponding to all the states in the 1D maze (Fig. 2B and 2C). At each time step, we update the firing rates of the nuclei in each module by evaluating Equations 1–13. The agent then compares the STN activity in the adjacent BG-modules and selects the direction corresponding to the module with higher activity.

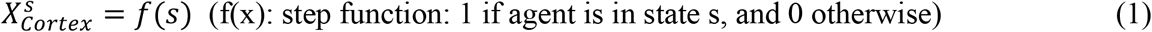

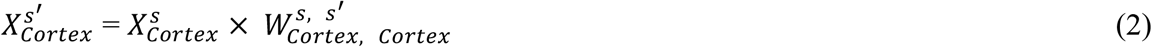

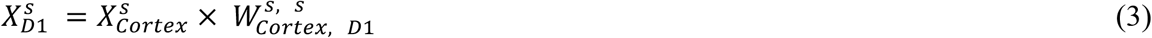

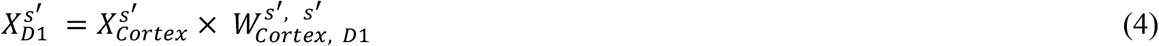

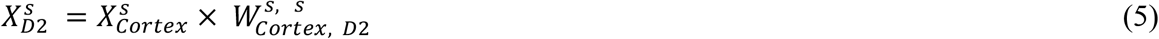

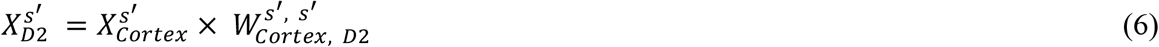

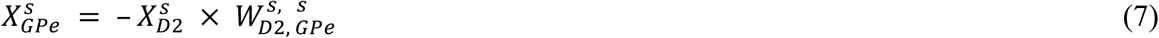

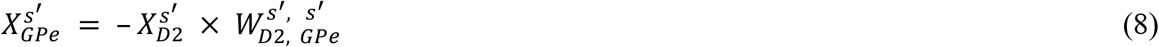

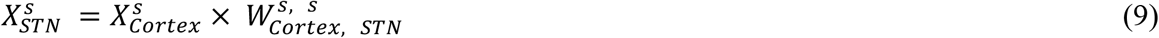

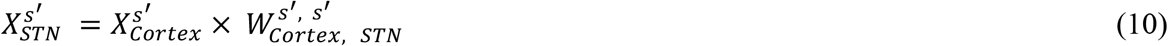

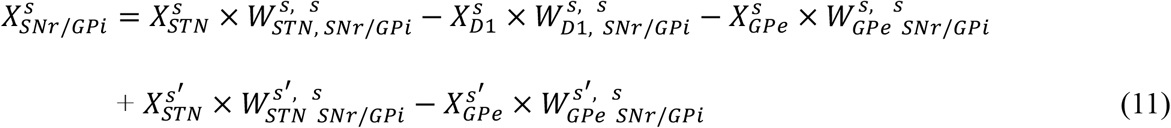

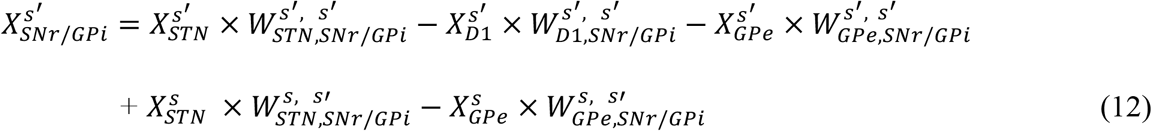

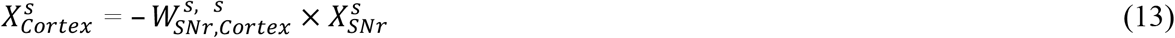

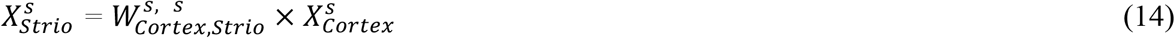

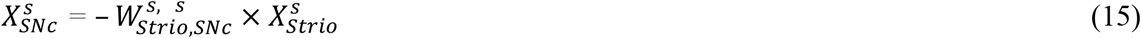

**Figure 2.**
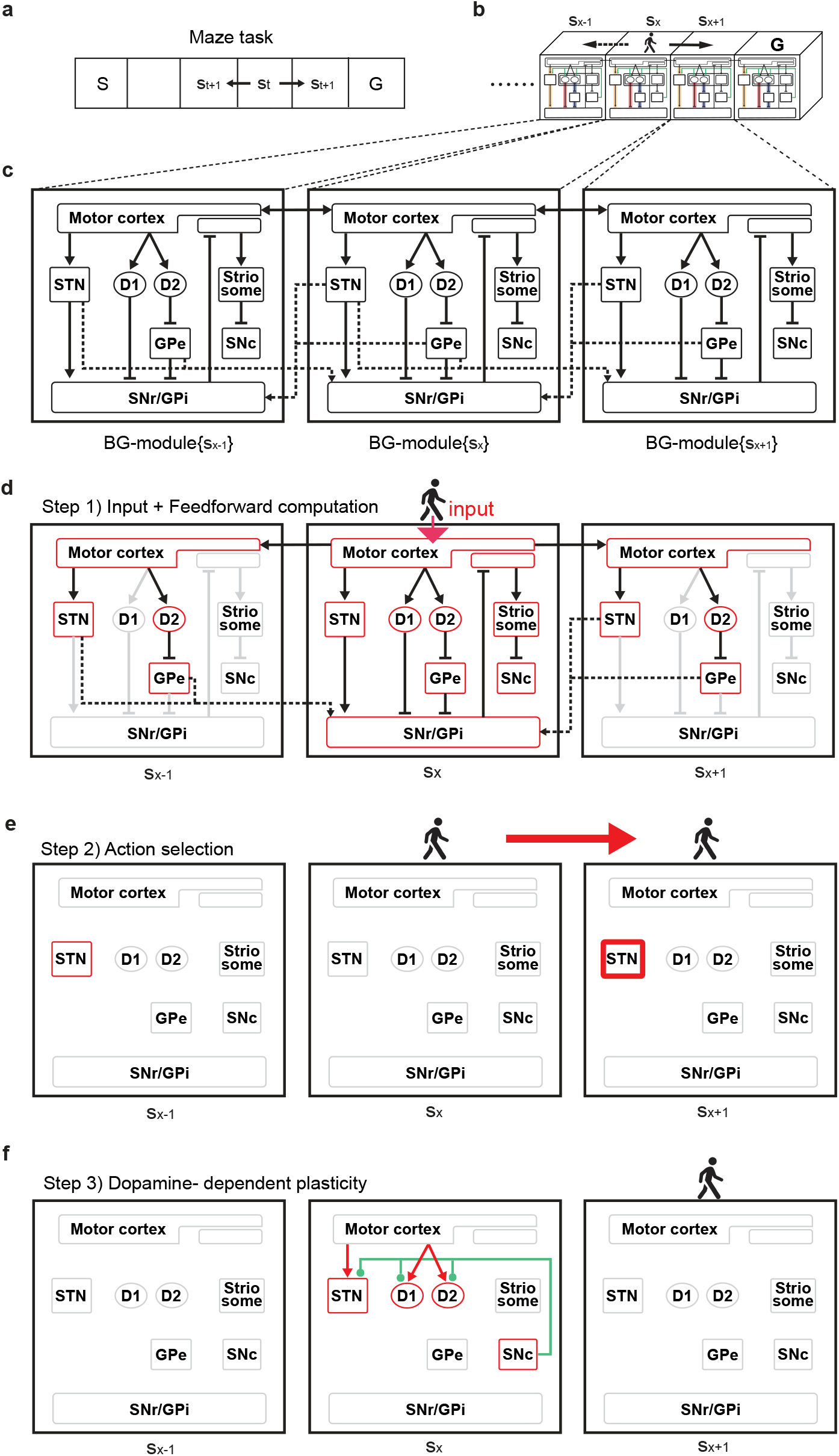
Tiled network moved through the maze by repeatedly selecting and learning actions. A. Definition of states in the maze. B. BG-modules were configured in a parallel manner, each module corresponding to each state. C. BG-modules were interconnected between transitionable states via inter-module projections. D–F. When an agent transitions to a new state, after a feed-forward computation (top), the next action is determined, the agent transitions to the next state (middle), and the weights are learned through dopamine release (bottom).

*X*_*cortex*_: cortical neurons; *X*_*D*1_: Striatal direct pathway neurons; *X*_*D*2_: Striatal indirect pathway neurons; *X*_*GPe*_: Globus pallidus external segment neurons; *X*_*STN*_: Subthalamic nucleus neurons; *X*_*SNr*_: Substantia nigra reticulata neurons; *X*_*SNc*_: Substantia nigra compacta neurons. Here, the superscript *s* represents the BG-module for the current state, and *s*^′^ represents the BG-modules for any possible next state. For example, if the current state *s* is *s*_*x*_, then *s*^′^ refers to *s*_*x-1*_ and *s*_*x+1*_. The superscript on the weight indicates a connected BG-module. For example, 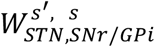 represents the weight from 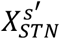 to 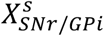, which is an inter-module connection.

When the agent enters a specific state (i.e., a specific place in the maze task), the cortex in the corresponding BG-module receives input from the environment (state observation; Equation 1). We set the model so that the cortical neurons in the module become active only when the agent is in a specific state. The activity of the cortex corresponding to the current state 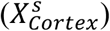 is then transmitted to the cortex in adjacent BG-modules 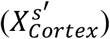 (Equation 2). Subsequently, feed-forward computation was performed for all nuclei of each adjacent BG-module (Fig. 2D; Equations 3–15). These computations were performed sequentially for each neuron using the firing rate model neurons. After a single update pass, the activities of the nuclei were fixed until the agent moved to the next state.

At the current state *s*, the agent determines the next state based on the activities of the STNs in the adjacent BG-modules 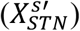, which correspond to the possible next states (left and right). The agent moved to a state in which the activity of the STN showed the maximum value among the two adjacent BG-modules (Fig. 2E). The state *s*_*t+1*_ was determined by the following equation, which is the ε-greedy method:

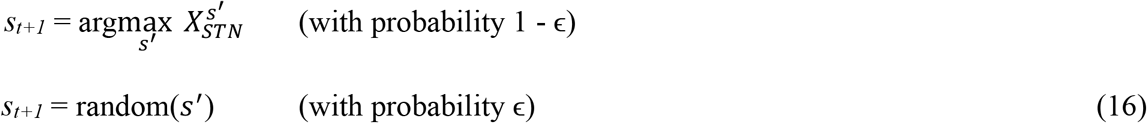

Then, the action *a*_*t*_, which causes the transition from *s*_*t*_ to *s*_*t+1*_, is performed. Note that our model is a state-value-based RL model in which we did not explicitly model action selection. Instead, actions are selected to enable the agent to transition to the next chosen state, which the detailed mechanism is described later. We next, however, our model to an action state model, in which the BG-modules are arranged across the action-state space. In this expanded model, each BG-module corresponds to a specific action within a given state, allowing the model to select an action by comparing the activity levels of the STNs in the modules associated with each potential action.

Finally, dopamine-dependent Hebbian plasticity was implemented to the cortex-striatum and cortex-STN connections (Fig. 2F). This rule is based on the biological evidence that dopamine influences synapse formation and that sub-second-level temporal coincidence of glutamate and dopamine release in the striatum facilitates synaptic plasticity^17,22^, which has been used in neural simulations^23^. Specifically, the weights were updated using the following equation:

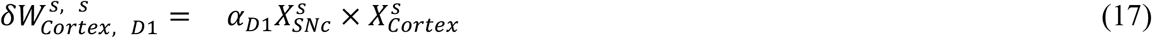

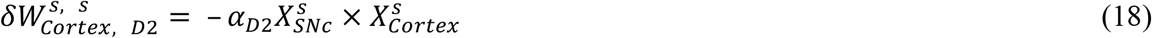

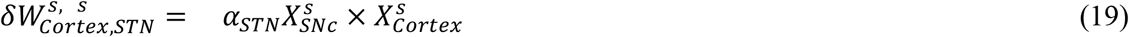

where 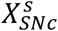 represents the activity of the SNc, corresponding to the amount of dopamine released in the striatum. For details on the update rates α_D1_, α_D2_, α_STN_, initial values of weights, and other relevant parameters are provided in Table 1 and 2.

**Table 1.** Initial values of the variables.

| Variable | Initial | Description |
| --- | --- | --- |
|  | Value |  |
| $X_{\text{Cortex}}$ | 0 (1.0) | Initial value of activity of cortex.<br>Cortex in the BG-module corresponding to the current state $s$ were assigned<br>an initial value of 1.0, then the rest of the cortex were set to the<br>initial values of 0. |
| $X_{\text{D1}}$ | 0 | Initial value of activity of direct pathway striatum (D1) |
| $X_{\text{D2}}$ | 0 | Initial value of activity of indirect pathway striatum (D2) |
| $X_{\text{GPe}}$ | 0 | Initial value of activity of external segment of globus pallidus (GPe) |
| $X_{\text{STN}}$ | 0 | Initial value of activity of subthalamic nucleus (STN) |
| $X_{\text{SNr/GPi}}$ | 0 | Initial value of activity of substantia nigra reticulata (SNr), and<br>Internal segment of globus pallidus (GPi). |
| $X_{\text{SNc}}$ | 0 | Initial value of activity of substantia nigra compacta (SNc) |
| $W_{\text{Cortex, D1}}$ | 1.0 | Initial value of weight between cortex to striatum (D1) |
| $W_{\text{Cortex, D2}}$ | 1.0 | Initial value of weight between cortex to striatum (D2) |
| $W_{\text{Cortex, STN}}$ | 1.0 | Initial value of weight between cortex to STN |

**Table 2.**
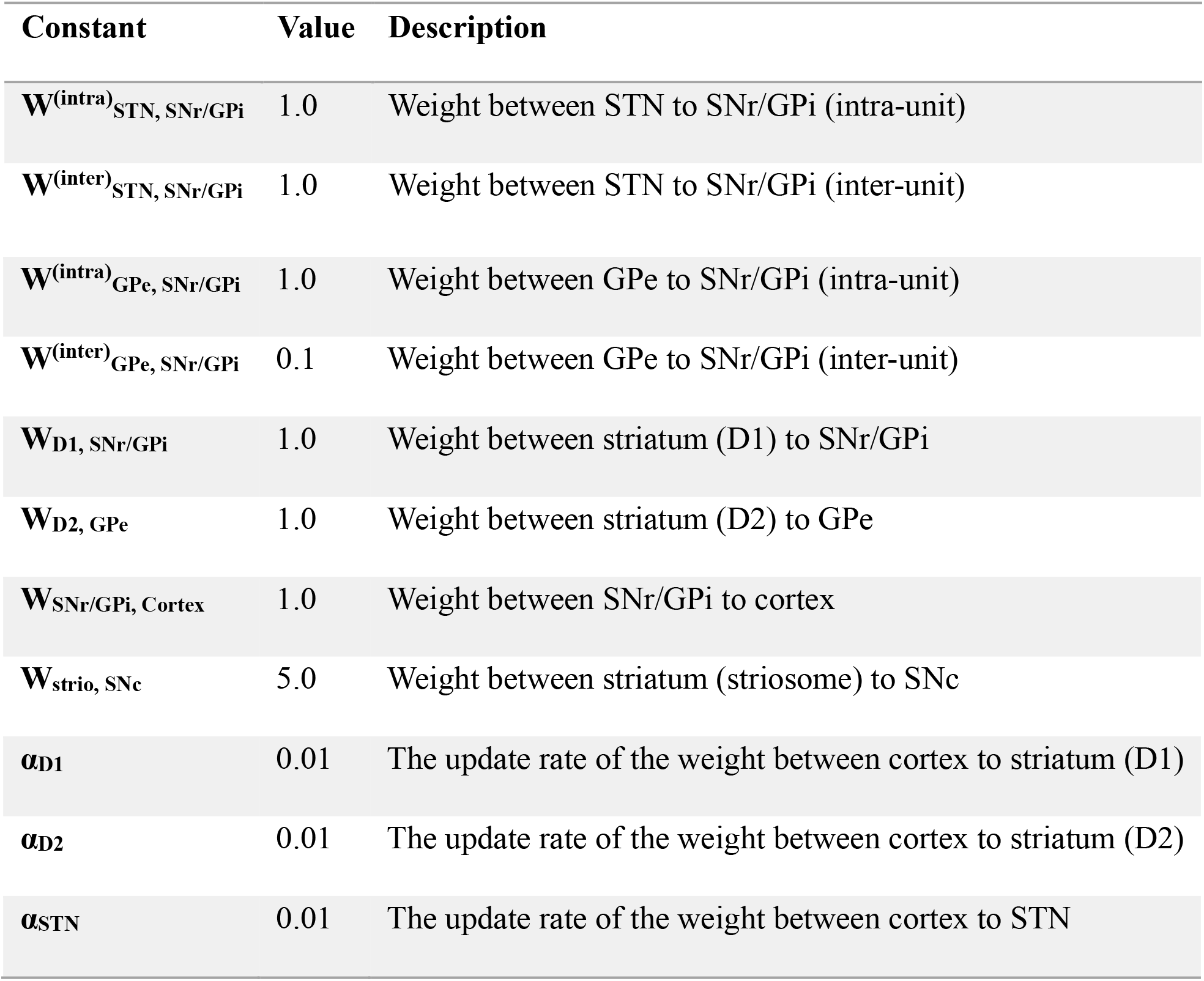
Constant value of the parameters.

### Tiled network can perform 1D maze task

We tested the learning capability of the tiled network using the 1D maze task. The period from the start to achieving the goal was defined as an episode, and a session was defined as having 200 episodes. As the training proceeded, the number of steps required to reach the goal decreased (Fig. 3A). For comparison, we also implemented a standard temporal difference (TD) learning approach using state value (commonly referred to as ‘V-learning’) as described in established RL frameworks^10^ (see details in Methods). The tiled network demonstrated a similar learning curve to V-learning (Fig. 3A). To understand the mechanism of the tiled network in performing the maze task, we investigated the dynamics of the network during the learning process.

**Figure 3.**
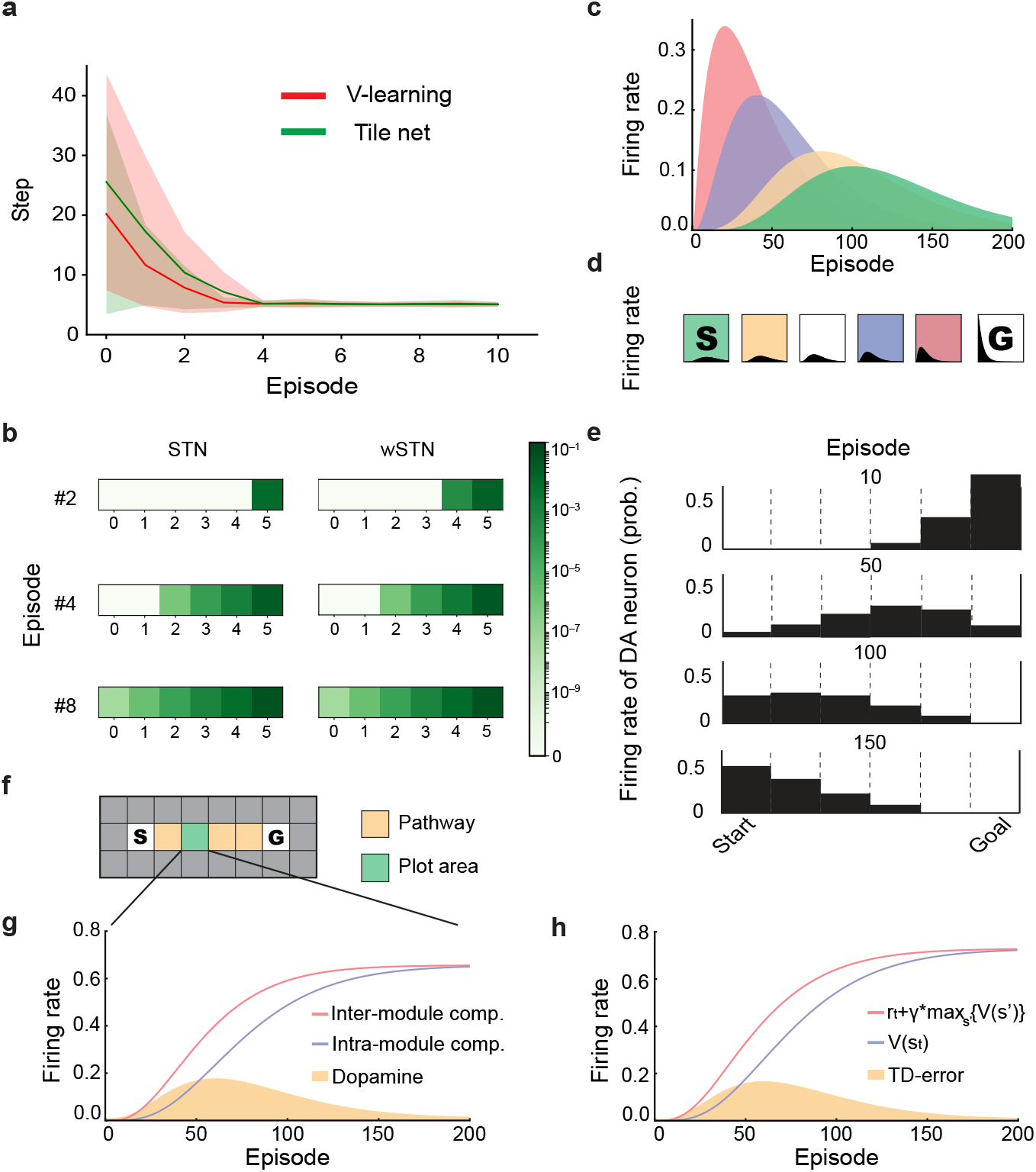
Tiled network learned 1D-maze task. A. Learning curves of the maze task with the state value learning model (red) and the proposed model (green). The solid lines represent the mean values and shades represent the range of mean ±1 SD (n = 100 trials per model). Vertical axis: number of steps; horizontal axis: number of episodes. B. Heatmaps of STN activity (top) and the synaptic weights from the Cortex to the STN (bottom) within each BG-module corresponding to individual states. From left to right, the heatmaps show activity levels (top) and weights (bottom), individually, at the 2th, 4th, and 8th sessions after learning began. C. Transition diagram of SNc activities within a session. Four exemplar BG-modules (red, blue, yellow, green) were selected in order of proximity to the goal, and the activity of SNc in each BG-module was illustrated over one session. Vertical axis: SNc activity; horizontal axis: episode counts. D. The diagram showing the transitions of SNc activity along the path learned by the agent. Vertical axis: SNc activity; horizontal axis, episode counts. E. Diagram showing the activity level of SNc from start to goal. Different learning stages (episodes 10, 50, 100, and 150) are shown. Vertical axis: SNc activity; horizontal axis: states from start to goal. F. Overview of the maze task. The state which the activity is displayed in (F) and (G) is highlighted by green shading. G. Transitions of the inputs to SNr/GPi through inter-module projection and intra-module projection during the maze task. The input through inter-module and intra-module projections in an exemplar BG-module is illustrated along with the session. Vertical axis: input; horizontal axis: number of episodes. H. Transitions of the values (γV_t+1_ and V_t_) during learning the maze task by the state value model. The values used to calculate the TD-error when the agent reached the same state as shown in (B) are illustrated. Vertical axis: values (γV_t+1_ and V_t_); horizontal axis: number of episodes.

In our tiled network, the agent moves in the direction in which the adjacent BG-modules have the highest STN activity. Thus, to move appropriately in the maze, the activity of the STN in the appropriate adjacent BG-module must be reinforced. The activity of the STN first increased at this goal (Fig. 3B, top). As the episodes progressed, this increased activity gradually shifted from the goal to the starting position (Fig. 3B, middle and bottom panels). Because the activity of the STN is modified by the plasticity between the cortex and the STN (Equation 18), the weight between the cortex and STN also showed the same propagation pattern; the weight 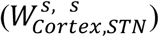 first became stronger near the goal, and as episodes progressed, it gradually propagated towards the start position (Fig. 3B). Given that the plasticity between the cortex and STN is driven by dopaminergic input from the SNc (Equation 18), the activity of the dopaminergic neurons in the SNc (*X*_*SNc*_) should also change accordingly. Indeed, the activity of the SNc first increased near the goal and then gradually propagated towards the starting position (Fig. 3C-E).

The mechanism of this propagation can be explained by the inter-module projection. Clearly, the trigger for this propagation was the reward given at the goal, which triggered dopaminergic activity in the BG-module in the goal state. As a result of the subsequent synaptic plasticity, the weight in the goal state 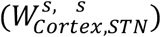 first became stronger. Notably, in the next episode, this increased STN activity in the goal state had an impact on the BG-module one step before the goal, since these modules have a connection via the inter-module projection 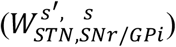. Because of this connection, increased STN activity in the goal state caused increased SNr/GPi activity one step before the goal (Equation 11), which would then lead to increased dopamine activity (Equations 13–15). This inter-module-driven dopamine transition reinforced the BG-module one step before the goal, which ultimately induced the propagation of the learning signal further backward to the start position in later episodes.

### Inter-module projection enabled tiled network to calculate TD-error

The tiled network as described functions as though it operates by a TD learning model in which learning is driven by temporal prediction errors. To explore this possibility, we compared neural activity in the tiled network with the computational process of V-learning. We analyzed strength of the input to the SNr/GPi by separating the input into inter- and intra-module components and examined their transitions across trials within a session (Fig. 3F). We trained the V-learning model using the same task, and compared the transitions of the state values (*Vs* and *Vs’*) to the intra- and inter-module inputs in the tiled network. The inter-module component initially showed an increase, followed by an increase in the intra-module component, which eventually matched the inter-module’s activity by the end of the session (Fig. 3G). The analysis demonstrated that the intra-module components mirrored the current state value *Vs*, whereas the intra-module components mirrored the next state value *Vs’* (Fig. 3H). Given that it is calculated by subtracting *Vs’* from *Vs*, the TD error can be calculated using two types of projections in our proposed network (inter-module and intra-module), enabling the network to update the state value.

### Tiled network can perform 2D maze task

We next extended the model to a 2D maze (Fig. 4A). As in the 1D maze, we arranged the BG-modules horizontally in a grid corresponding to the spatial layout of the maze (Fig. 4B). In a 2D maze, there are four possible next states (up, down, left, and right). Therefore, the BG-module in the current state receives inter-module projections from the four BG-modules corresponding to all possible subsequent states (Fig. 4C and 4D). The tiled network successfully learned the 2D maze and performed comparably well with TD learning (Fig. 4E). During learning, dopamine activity propagated from the goal to the starting point, as observed in the 1D maze (Figs. 4F and 4G).

**Figure 4.**
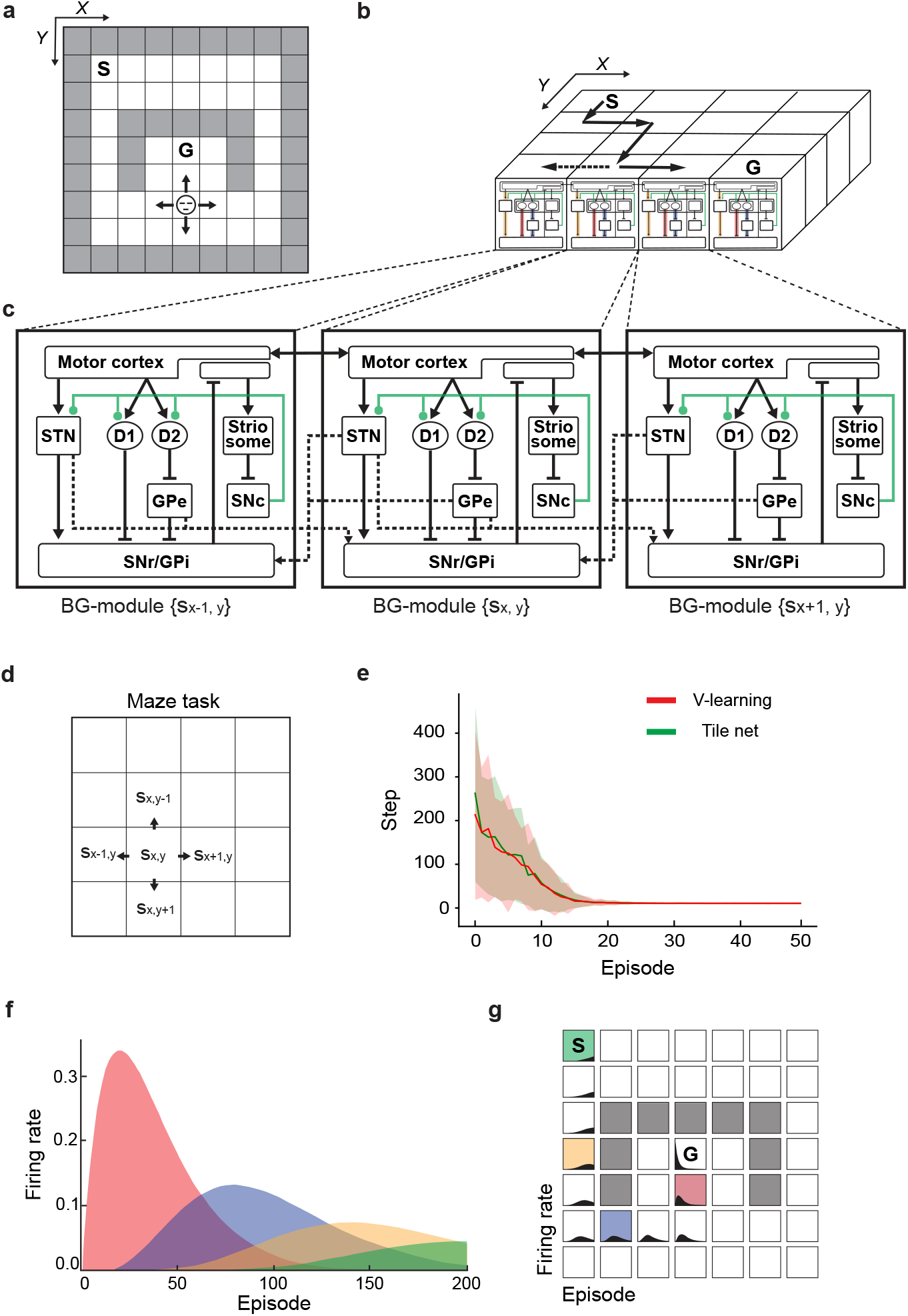
Tile Net can be extended to two-dimensional mazes. A. Overview of the maze task, a 9×9 maze with obstacles. B. BG-modules are placed in parallel on the two-dimensional maze. C. Each BG-module is connected by an Inter-module projection only between transitionable states. D. Agents can move up, down, left, and right. E. The tile net can also learn a two-dimensional maze. F and G. Dopamine activity (SNc) during learning transitioned from the goal to the start, as in the 1D maze.

In summary, our proposed model can perform RL by employing inter-module projections to propose a gradual propagation of the TD error, which is represented by dopamine activity. Ultimately, this propagation triggers corticostriatal plasticity, strengthening intra-module projections. As a result, dopamine activity decreases over time, forming bump-like dopamine signaling patterns.

### The balance between two inter-module projections represented discount rate

We demonstrated above that the subtraction between the intra-module and inter-module projections can represent the TD error in RL. We now demonstrate that the balance among the two inter-module projections 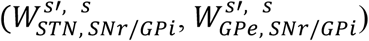 and the three intra-module projections 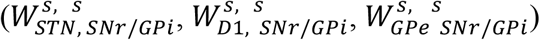, can determine the discount rate. The discount rate scales the contribution of future rewards to the TD error calculation using the following equation (see Satton and Barto (2018) for details):

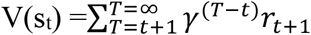

The higher the discount rate, the more future rewards are valued. In the proposed model, SNc activity, confirmed to be equivalent to the TD error, was calculated using the following equation:

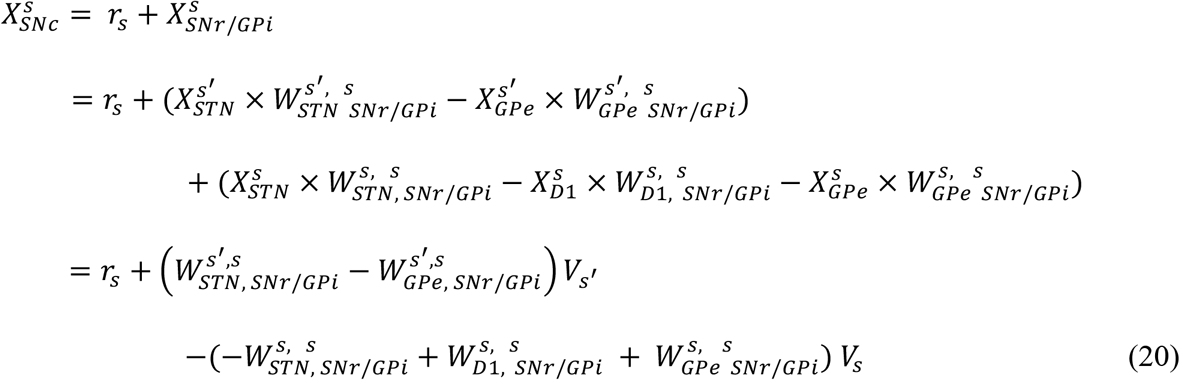

Here, we made two simplifications. First, we assume that 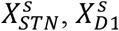, and 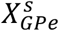 represent the state value of the current state, *V*_*s*_ (the 3^rd^ term in Equation 20). Second, we assume that both 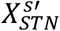 and 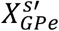 represent the state value of the next possible state, 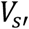 (the 2^nd^ term in Equation 20).

The activity of the STN was interpreted as the state value 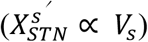 because the agent determines its movement direction based on the STN activities of adjacent BG-modules. Notably, both cortex-STN and cortex-D2 synapses were modified by dopamine-dependent plasticity to the same extent because their learning parameters were set to be equal (Equations 17 and 18; see Table 1). As a result, when the cortex-STN synapses were strengthened, the cortex-D2 synapses were weakened at an equivalent rate. Therefore, if STN activity represents the state value, then D2 activity represents the negative state value 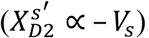. Given that the GPe receives inhibitory input from D2 neurons, GPe activity can therefore represent the state value 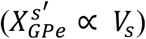. We then defined the term *γ* ^(inter/intra)^ as follows:

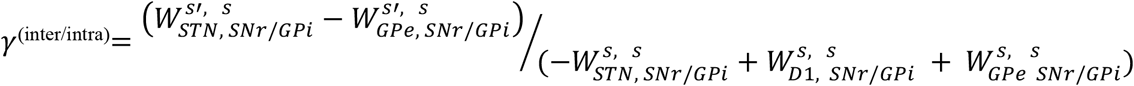

Substituting this definition into Equation (20), we obtained:

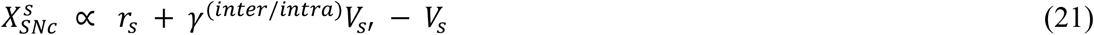

This is the definition of the TD error calculations. Thus, we hypothesized that *γ*^(inter/intra)^, which is the balance of the two inter-module and three intra-module projection weights, functions as a discount rate used in RL.

To test this hypothesis, we used a conflict task, a maze task with two goal states with different reward values: a large reward located at a long distance and a small reward located at a short distance. Specifically, we set up a high-cost-high-return reward with a long timeline to receive a large reward (cost: six steps; reward: +3) and a low-cost-low-return reward with a short timeline to receive a small reward (cost: five steps; reward: +1) (Fig. 5A).

**Figure 5.**
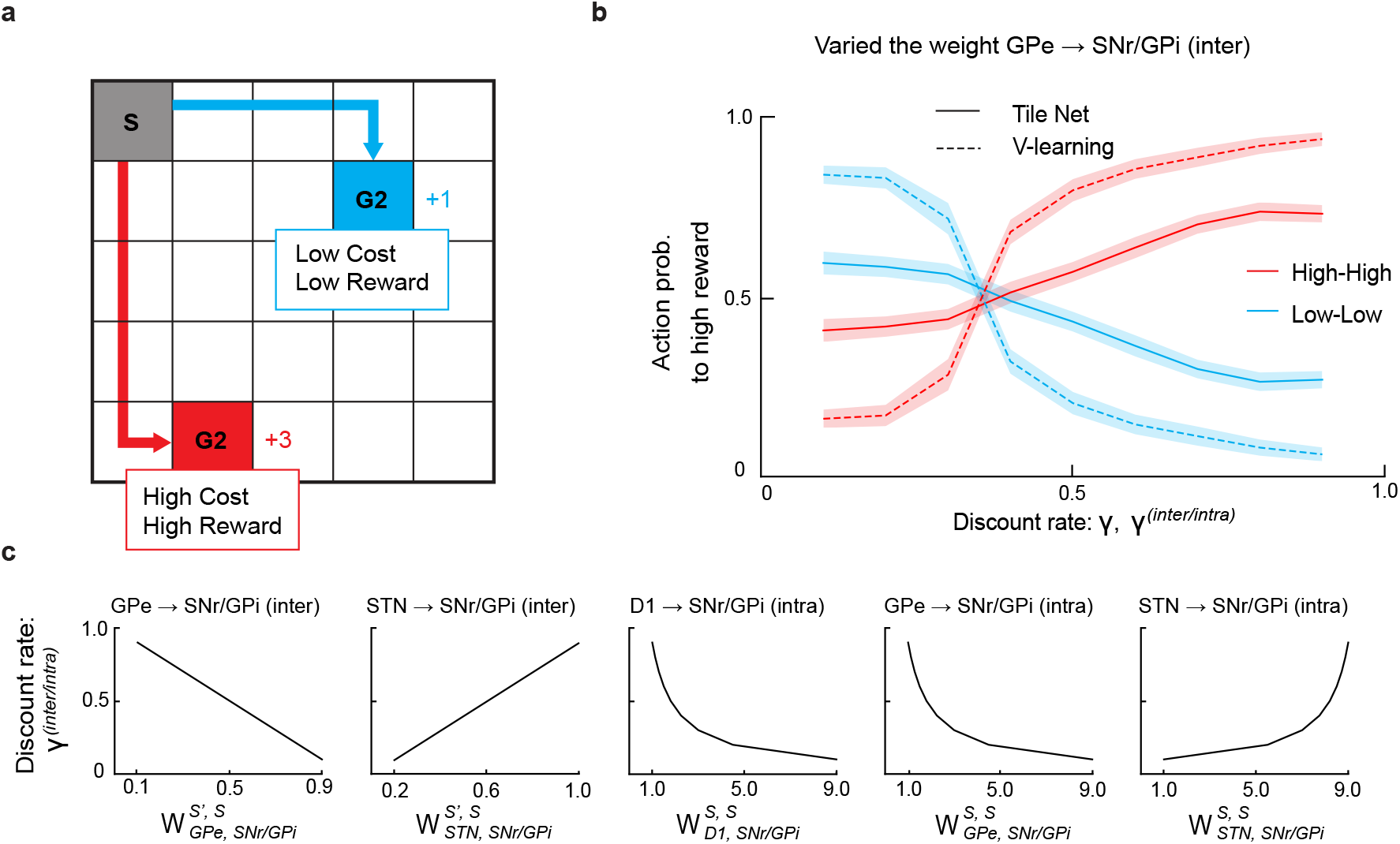
The strength of inter-module projections in the tiled network determined the discount rate. A. Overview of the conflict task, which includes two types of rewards: a “low-cost, low-return reward” with a short distance to the reward but a smaller reward, and a “high-cost, high-return reward” with a longer distance to the reward but a larger reward. B. The probability of selecting each reward type as a function of the inter-module projection strength (γ^(inter/intra)^) or discount rate (γ). Both the tiled network and state-value TD-learning models were trained on the conflict task for 200 sessions, with γ^(inter/intra)^ and γ ranging from 0.1 to 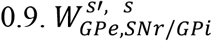 was modified to make *γ*^(inter/intra)^ range between 0.1 and 0.9, while all the other four weights were held constant. For the tiled network, solid lines represent the average selection probabilities for the high-cost-high-return reward (red) and low-cost-low-return reward (blue), with shading indicating the ±1 standard deviation (SD) range. A comparison with state-value TD-learning (V-learning) is shown, where dashed lines represent the selection probabilities. Vertical axis: probability of selecting each reward; horizontal axis: the projection strength or discount rate (γ^(inter/intra)^ for the tiled network and γ for TD-learning). Shading indicates the ±1 SD range for both models (red: high-cost-high-return; blue: low-cost-low-return). C. Changes in γ^(inter/intra)^ as a function of each weight. Each projection weight 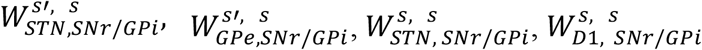, and 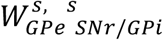 was individually varied to set γ^(inter/intra)^ across a range from 0.1 to 0.9.

We tested how the agent behaves in the conflict task by varying γ^(inter/intra)^ through the manipulation of individual projection weights, including 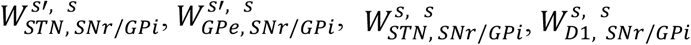, and 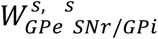. The agent tended to choose the low-cost, low-return reward under conditions where γ^(inter/intra)^ was low. As the value of γ^(inter/intra)^ gradually increased, the probability of the agent choosing the high-cost, high-return reward increased (Fig. 5B). The observed behavioral change can be understood in that, as γ^(inter/intra)^ increased the agent came to place greater importance on future rewards and learned to maximize cumulative rewards over a longer term. Considering the possibility that γ^(inter/intra)^ may change systematically by varying the individual projection weights (Fig. 5C), this result suggests that the balance of projection strengths among those five pathways was a parameter that modified the discount rate.

In addition, we compared the influence of the discount rate represented in the conventional V-learning model with that represented in the proposed model. We systematically varied the discount rate γ in the V-learning model from 0.1 to 0.9, matching the range used for γ^(inter/intra)^ in the tiled network. For each value of γ, we measured the probability of the agent selecting either the high-cost-high-return reward or the low-cost-low-return reward. The V-learning model exhibits a similar trend, with higher discount rates leading to a greater likelihood of choosing a high-cost, high-return reward (Fig. 5B). These results suggest that the balance of the strengths between the two inter-module projections represents the discount rate.

### Tiled networks can be extended to action value models

We introduced a biological RL model in which each BG-module corresponds to a state and demonstrated that it functions similarly to the conventional TD-learning model using state values, a V-learning model. In Q-learning, another learning method, the cumulative expected future reward for taking a specific action in each state is defined as the action value, also known as the Q-value, often denoted as Q(*s, a*). In the Q-learning model, the agent learns the optimal action strategy by updating the Q-value^10^.

We introduced an action value model of the tiled network by aligning BG-modules across the entire state-action space (*s, a*) ∈ ***S*** × ***A***, such that each BG-module corresponds to a unique state-action pair (Fig. 6A and 6B; see details in Methods). This configuration ensured that every possible combination of state and action had a corresponding BG-module, enabling the model to learn and update the Q-values for each state-action pair.

**Figure 6.**
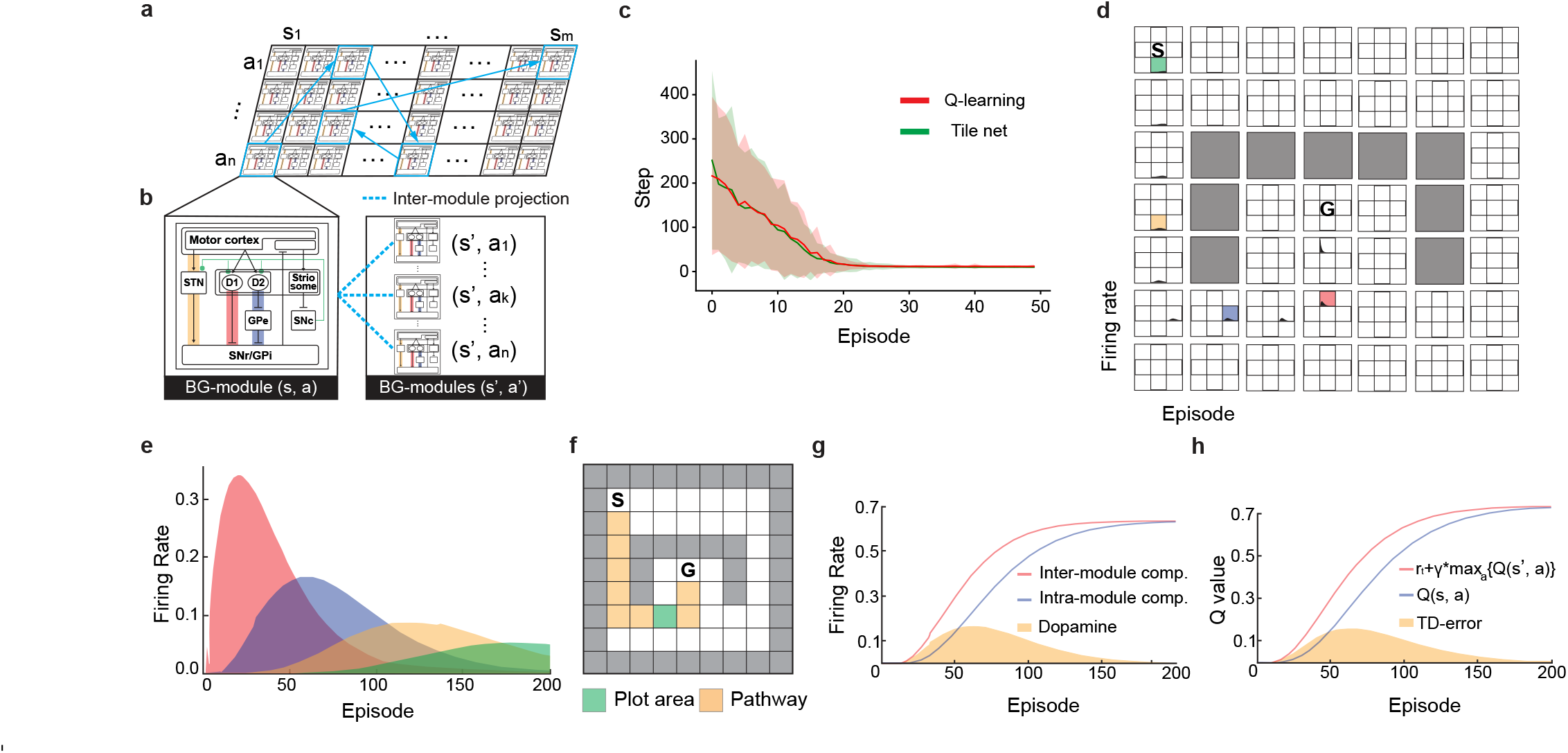
Tiled network was capable of learning action values. A. Tiling the BG-modules on the state-action space. B. Schematic of inter-module connections in the action value model. Inter-module projections exist between the modules corresponding to (*s, a*) and all the modules corresponding to (*s*′, *a*_*1*_), (*s*′, *a*_*2*_), …, (*s*′, *a*_*n*_). Here, *s* is the current state, a is the action taken in state *s, s*′ is the next state, and *a*_*1*_ to *a*_*n*_ are the possible actions in the next state. C. Learning curves for the maze task using Q-learning (red), state value tiled network (blue), and action value tiled network (green). The solid lines represent the mean values, and the shading represents the range of mean ±1 SD, calculated across 100 sessions. Vertical axis: number of steps required to reach to the goal; horizontal axis: number of episodes. D. Transition diagram of SNc activity within a session. Four exemplar BG-modules (red, blue, yellow, green) were illustrated, showing SNc activity over one session. Vertical axis: SNc activity; horizontal axis: number of episodes. E. Transition of SNc activity along with the learned path. The four exemplar BG-modules highlighted in (D) are emphasized with the same color shadings. Vertical axis: SNc activity; horizontal axis: number of episodes. F. Overview of the maze task. Modules corresponding to the state-action pairs with displayed activity in (D) and (E) are highlighted with a green grid. G. Transition of the inter-module and intra-module projections to the SNr/GPi during the maze task. Inputs through inter-module and intra-module projections in an exemplar BG-module were shown over the session. Vertical axis: input ; horizontal axis: number of episodes. H. Transition of action values (γQ_t+1_ and Q_t_) during the maze task learning with the Q-learning model. The TD-errors when the agent reached the same state as in (G) are shown. Vertical axis: values (γQ_t+1_ and Q_t_); horizontal axis: number of episodes.

We then performed learning in the 2D maze task using the state-action configuration of the BG-modules. When the agent moved to a particular state, cortical inputs were provided to the four BG-modules, each corresponding to one of the possible actions available in that state. Feed-forward computations were then performed for each module (See Equations 25–39 in the Methods section for further details). The agent then selected the action corresponding to the BG-module with the highest STN activity. Regarding inter-module projections, we introduced a gating mechanism that allowed only the projection from the adjacent BG-module corresponding to the highest possible action in the next states (i.e., the actions with the highest STN activity in the next states) to influence the SNr/GPi of the current state (Equation 35 in the Methods). This means that only those winning in the next state can project to the BG-module of the current state-action pair. We also used dopamine-dependent Hebbian plasticity for weight updates.

Using the above settings, we performed the maze task. For comparison, we also implemented a standard conventional Q-learning model^10^ (see the detailed implementation in Methods). Our proposed action value model could learn the maze task, performing comparably well to the standard Q-learning model (Fig. 6C). Additionally, we observed that the dopamine signals in the model propagated backward from the goal to the start, similar to the state-value model (Fig. 6D and 6E). Furthermore, we analyzed dopamine activity in the tiled network and compared it with the TD error calculated in the Q-learning model. We found that the dopamine signal in our model arose from the difference between the inter-module projections (which represent the maximum expected future reward among the next state-action pairs) and the intra-module projections (which represent the current state-action value). This difference effectively computes the TD error, similar to the results obtained using the state-value model (Fig. 6F and 6G). Based on these results, we demonstrate that the action value model performs computations analogous to those of conventional Q-learning, suggesting that our biological model captures the essential mechanism of Q-learning.

### Action value-based tiled network can perform cart-pole task

To evaluate the generalizability of the proposed model across various tasks, we used the action value model of the tiled network and trained it on a cart-pole task. During the cart-pole task, the agent controlled a cart that could be pushed to the left or right to balance a pole hinged on its top (Fig. 7A). The goal was to prevent the pole from falling over by applying force to the cart and keeping the pole upright for as long as possible. The task ended when the pole fall beyond a certain angle or when the cart moves beyond the boundaries of the track. In this task, the agent can observe the following four state variables at each step: 1) position of the cart, 2) velocity of the cart, 3) angle of the pole, and 4) velocity of the pole tip. Although these four variables were continuous values, we divided the values of each variable into six bins, resulting in 6^4^ = 1296 possible states. At each step, the agent can take one of two actions: applying fixed power (acceleration) to the cart, either to the right or left. Because power is constant, there are only two possible actions. The maze task had only four states, with the next state uniquely determined once the state *s*_*t*_ and action *a*_*t*_ were determined. In the cart-pole task, however, the number of next states was 1296, and the next state was not uniquely determined even with the same pair of *s*_*t*_ and *a*_*t*_ values due to the discretization of the continuous variables. Small differences in the continuous state variables can lead to different discretized next states, even when starting from the same discretized state and performing the same action, making it more challenging than the maze task.

**Figure 7.**
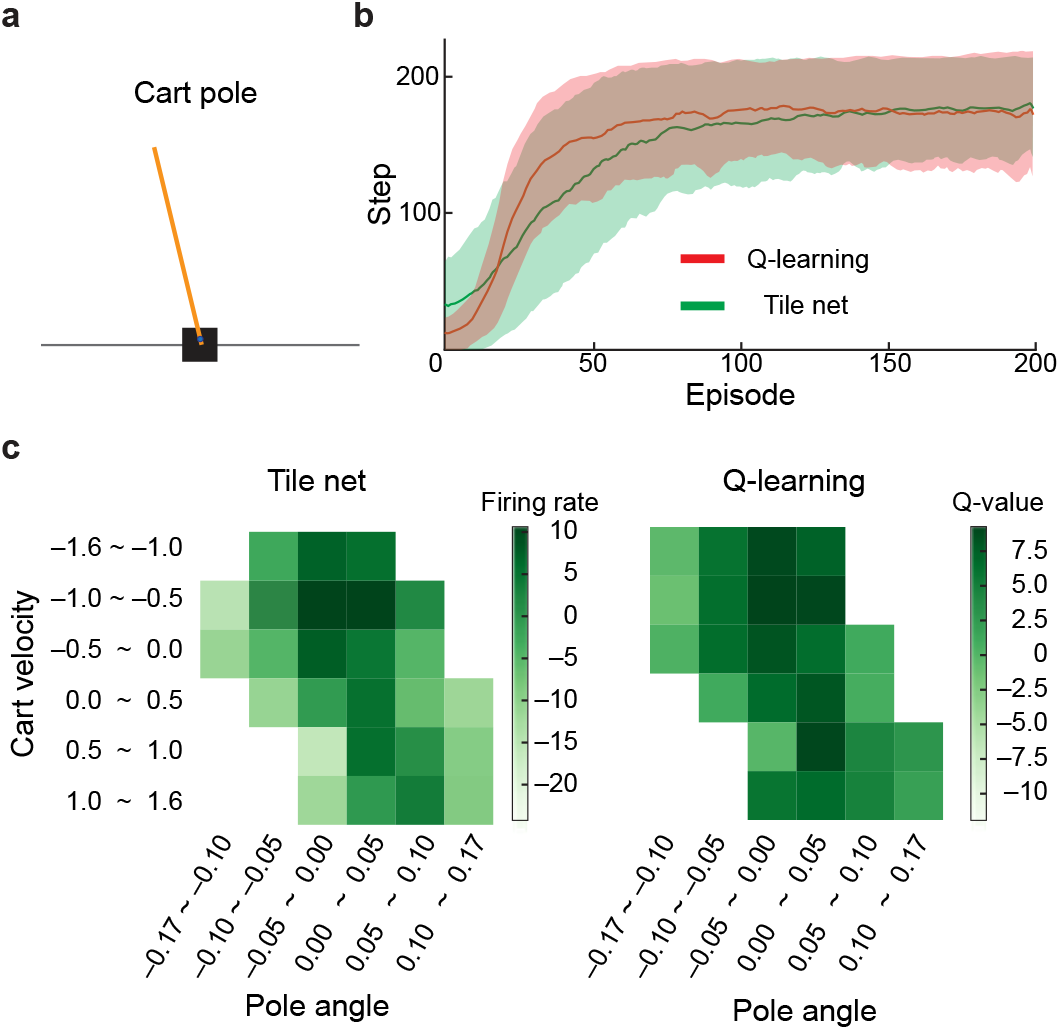
Tiled network learned the cart-pole task. (A) Overview of the cart-pole task. The cart-pole task involves keeping a pole balanced on a cart as long as possible without it falling over. (B) Learning curves for the cart-pole task. The learning curves of the action value tiled network (green) and Q-learning (red) are illustrated. The learning curve for each episode indicates the number of steps the pole was kept balanced without falling. Each model was trained for 200 sessions, and the average number of steps is shown with solid lines, with the ±1 SD range indicated by shading. Vertical axis: number of steps; horizontal axis: number of episodes within the session. (C) Heatmap of the STN activity after learning the cart-pole task. The activities of the STN within the BG-module corresponding to each state-action pair (left), as well as the Q-values from Q-learning (right), are shown as heatmaps. For the STN activities, the highest activity among the BG-modules corresponding to multiple state-action pairs in each state was illustrated. Of the four states of the cart pole, the 2D heatmaps illustrate two dimensions (cart position and pole angle). The STN activities and Q-values were averaged across the remaining dimensions for visualization purposes.

In this model, each BG-module represents a specific state-action pair. When the agent resulted in a specific state, *s*, cortical inputs were provided to the two BG-modules corresponding to (*s, a*_*left*_) and (*s, a*_*right*_), representing two possible actions in that state. After sequentially computing the firing rates of the nuclei in each BG-module (Equations 25-37–Methods), the action corresponding to the BG-module with the highest STN activity was selected. Subsequently, the weights were updated using dopamine-dependent plasticity. The activity levels are reset before transitioning to the next state. This cycle was repeated until the cart-pole task was complete.

We found that the tiled network quickly learned the cart-pole task, increasing the number of steps per episode from approximately 20 to more than 195, similar to the Q-learning model (Fig. 7B). This result indicates that the tiled network is a capable RL model that can learn complex tasks such as cart poles, demonstrating its potential applicability to a range of RL tasks. We calculated the STN activities in the BG-modules corresponding to each state-action pair (*s*_1_, *s*_2_, *s*_3_, *s*_4_, a) ∈ (***S***^4^ × ***A***). We found that the STN activity was higher in states near the center position and where the pole angle was close to zero (Fig. 7C, left). For visualization, we displayed a heatmap of STN activity on a two-dimensional plane, focusing on two axes: *s*_1_ (cart velocity) and *s*_2_ (pole angle), averaged over *s*_3_ and *s*_4_. For comparison, we generated an action value map by plotting the learned Q-values using the conventional Q-learning model for the same state-action pairs on the *s*_1_ and *s*_2_ axes. The similarity between the heat maps of the STN activity (Fig. 7C, left) and Q-learning action values (Fig. 7C, right) indicates that the STN activity in our model effectively represents the learned action values.

Based on these results, our proposed tiled network demonstrates the capability of learning Q-values using the TD-error method in tasks such as the cart pole, suggesting its potential applicability to other RL tasks where states and actions can be represented in a tabular form.

## Discussion

In this study, we propose a biologically inspired RL model, the tiled network, which incorporates key anatomical features of the basal ganglia. The proposed model includes modular cortical-basal ganglia circuits with hyperdirect, direct, and indirect pathways having different projection ranges. By tiling these modules horizontally, we demonstrated that the model can perform RL. Notably, the ability of the model to compute the TD error arises from differences in the projection ranges between parallel circuits, which are physiologically well documented. Based on these results, we hypothesize that RL in the basal ganglia is achieved through the computation of values at the modular level and the propagation of these values through inter-module connections.

The dopamine signal propagation observed in our proposed model aligns with the neural activity patterns documented in the dopamine system of animals. Non-human primate studies investigating associative learning between sounds and rewards have reported similar dopamine propagation, in which the activity of dopamine neurons initially increased after reward delivery; however, over time, they no longer responded to the reward itself and started responding to the associated sounds^2,24^. In addition, a recent experiment using rats demonstrated that during the maze learning task, dopamine signals propagated in a bump-like pattern from the goal back to the start position^25^, closely matching the predictions of the TD-learning model. Consistent with these findings, our results demonstrate that dopamine activity initially peaks at the goal position and gradually shifts backward to earlier states over time (but note that some studies suggest this is not always the case with respect to striatal dopamine release^26^). This finding supports the existing research hypothesis that dopamine encodes reward prediction errors by calculating the TD-error. This process in our model arises from inter-module projections, which govern the information flow between modules and act as the discount rate in a RL context. Based on these results, we posit that our biologically inspired model could reflect the mechanism of dopamine signaling as it has been observed.

In this model, RL was executed by tiling the modules of the basal ganglia into individual columns based on several anatomical and physiological facts. Similar to the cerebral cortex, the striatum, STN, and SNr have topographical specializations, such as those responsible for specific body parts^4,5^. This indicates that the cerebral cortex and basal ganglia are organized into modules based on their topographies. In the striatum, these modules are the matrisomes that tile the large matrix compartment within which the smaller striosome compartment, also clustered into modules, resides^27^. Therefore, cortico-basal ganglia circuits may be arranged in a horizontally tilted configuration.

The basal ganglia receive projections from multiple cortical regions and process them through multiple parallel circuits. These circuits form separate loops for processing inputs from distinct cortical regions, such as the motor, oculomotor, prefrontal, and limbic areas^28^. In this context, models that integrate the motor and cognitive loops have also been proposed^29^. Our model is based on more recent findings, suggesting that basal ganglia circuits are further subdivided into detailed modules^30^. We investigated how these finer-scale modules perform information processing when integrating information across them.

We introduced dopamine-dependent Hebbian plasticity for weight updating. This rule is a modified version of the Hebbian rule, proposed as a synaptic strengthening system between pre- and post-synaptic neurons^31^. We introduced this mechanism based on the fact that the neural activity in the striatum, the input nucleus of the basal ganglia, is modulated by dopamine^32,33^. Dopamine-dependent Hebbian plasticity is triggered by dopamine input, indicating that the spatial and temporal regulation of dopamine input is crucial for performance. In our model, the activity of the SNc, corresponding to dopamine release, was directly controlled by striosome activity. This is based on the fact that striosomes can directly inhibit dopaminergic neurons^20,34^, making our model biologically plausible.

In RL, the characteristics of the agent’s behavior are determined by various hyperparameters such as the discount rate (γ). The neural mechanisms involved in the adjustment of such parameters in RL have been explained by the release of neuromodulators. For example, serotonin (5-HT) release has been hypothesized to encode a discount rate^35–37^. In our model, we attributed the discount rate (γ) to the balance among inter- and intra-module projection across the hyperdirect, direct, and indirect pathways. This hypothesis suggests that the balance of strength between different parallel circuits within the basal ganglia is crucial for appropriate behavior and learning. It could serve as a novel model for understanding impulsive behavior disorders, such as obsessive-compulsive disorder and attention deficit/hyperactivity disorder, in which dysregulation of basal ganglia circuits is implicated. Further work is required to integrate our hypothesis that the hyperparameters of learning are determined by the balance of strengths among multiple parallel circuits, with existing hypotheses that attribute these parameters to neurotransmitter release.

Our model has limitations. It requires the tiling of BG-modules across all state-action spaces, which may limit its ability to handle more complex tasks. To address this scalability issue, future studies can incorporate representation learning or function approximation to manage the exponential increase in the number of BG-modules required. Additionally, we used a simple firing-rate neuron model rather than a more biologically precise model, such as spiking neurons, which compromises its biological accuracy to some extent. Neural circuits that use spiking neurons are expected to exhibit different dynamics. Thus, whether the proposed model can perform RL using spiking neurons must be verified. In our model, the striosome contained neurons that inhibited the SNc. Striosomes contain both D1R- and D2R-expressing neurons that can influence dopamine neurons^38^; however, these details were not included in our current model. The similarity of the modules proposed here to the matrisomes based on corticostriatal and striatopallidal anatomical studies remains to be determined. To more accurately reflect the actual dopamine-modulating circuitry, future updates to the proposed model should incorporate the anatomical and functional insight.

The most striking feature of the proposed model architecture is that the modules, which are the units of calculation, are tilted horizontally. Each module consisted of a complete set of canonical circuits in the basal ganglia. These modules are interconnected by pathways between temporally proximal states, the states between which the agent can directly transition, leading to sequential activation. This design allows each module to perform its calculations and transmit its results horizontally, thereby achieving RL. Each module corresponds one-to-one with a state or an action-state pair, which is the core unit of RL. The individual responsibilities of module functions mirror the fact that each area of the brain has its own responsibilities. The architecture and function of the horizontally tiled network of modular calculation units are fundamentally different from those of artificial neural networks currently in use, which are based on multiple randomly connected layers with no individual responsibility. The horizontally tiled calculation module shows potential for new brain-inspired artificial neural network calculations, and it is worth pursuing further computational possibilities in the future.

## Methods

### Model summary

We implemented the cortico-basal ganglia network as a tiled network of modular circuits in which each module (BG-module) represented either a state or a state-action pair. Each BG-module consisted of the major nuclei in the basal ganglia: cortex, striatal-D1, striatal-D2, STN, GPe, SNr/GPi, striosome, and SNc. Each nucleus within the BG-module was implemented as a single firing rate neuron representing the average activity of that nucleus.

Each nucleus is connected by projections (weights). Most connections were intra-modular, representing pathways within the same BG-module. However, key inter-module projections were also included for the hyperdirect (STN) and indirect (GPe) pathways to the SNr/GPi projection. The initial values of the weights are listed in Table 1.

The agent interacts with the environment by performing actions to determine the next state. Upon entering a new state, the network performs feedforward computations through all nuclei within the corresponding BG-module as well as adjacent modules via inter-module projections. These computations, described by Equations 1–13, are executed in serial order. Once the calculations were complete, the firing rates of all nuclei were fixed until the next action was performed. The next action is selected based on the activity of the STN, and the synaptic weights are updated using dopamine-dependent Hebbian plasticity rules. Following this action, the network resets the firing rates and recalculates the feed-forward activations for the new state.

### Weight update

For weight modification, we employed dopamine-dependent Hebbian plasticity, inspired by the results of Yagishita et al. (2014)^17^. This plasticity rule was adapted for use in a firing-rate model to suit the averaged neural activity framework. Weight updates were performed according to Equations 17–19 after each feed-forward computation cycle. Modifiable weights included the cortex-striatal and cortex-STN connections, whereas all other weights were fixed throughout the simulations. The weights were preserved across episodes within a session and reset to their initial values at the end of each session.

### The 1D maze task

The maze consisted of six states arranged. The agent explored the maze, starting at the leftmost start point (S) and aiming to reach the rightmost goal (G) (Fig. 2A). The agent can move either to the left or to the right in each step. Upon reaching the goal, the agent received a reward of 1 and returned to the starting position. During the task, the period from the start to the goal was defined as one episode, and each action was defined as one step. If the agent bumped into a wall or obstacle, we counted that action as one step, and the agent restarted from the position before the collision. There were no collision penalties. In our maze task, we did not introduce a penalty for the action. The learning process of the tiled network in the maze task followed the procedure outlined below.

1. The agent started at the left-most state (S).
2. An activity of 1 was applied to the cortical neuron in the BG-module, which denotes current state (Equation 1).
3. Feed-forward computations were performed. The activities of the neurons in the BG-module of the current state s and the BG-modules of adjacent states s’ were then calculated (Equations 2–15).
4. The activities of the STN in the adjacent BG-modules are then compared, and the agent selects an action to move to the state with the highest STN activity (Equation 16).
5. Modifiable weights of the current state were then updated (Equations 17–19).
6. Activity levels across all BG-modules were then reset, and the agent transitioned to the new state.
7. Steps 2–5 were repeated until the agent reached the goal (end of an episode) or exceeded the limit of 200 actions.

An episode was defined as the period from when the agent started at (S) until it either reached the goal (G) or exceeded the limit of 200 actions. Each session comprised 200 episodes, during which the weights were not reset. At the end of each session, the weights of all BG-modules were initialized for relearning. During action selection, because this is a state-value-based model, the network can only determine the adjacent state with the highest value. Therefore, we employed an epsilon-greedy policy to determine the action of the agent (epsilon was set to 0.1).

To evaluate the tiled network, we used a temporal difference learning model with the state value (V-learning) and compared the results. Both models were trained for over 200 sessions, and the average number of actions required to reach the goal in each episode was calculated.

The standard deviation (SD) across all sessions was also calculated, and the range of ±1 SD was indicated with shading.

### Adjustment term for reward computation in SNc

Learning progresses by propagating the value of the reward based on the plasticity between the cortex and striatum, which is modified by dopaminergic input from the SNc. Here, the computations of SNr/GPi and SNc are given by Equations 22 and 23, respectively, which demonstrate that when SNr/GPi becomes active, SNc also increases activity. Therefore, without adjustments, SNc activity would disproportionately reinforce all states (or state-action pairs), including suboptimal states.

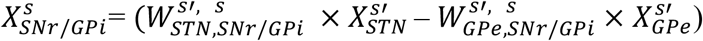

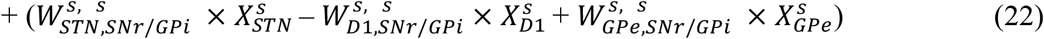

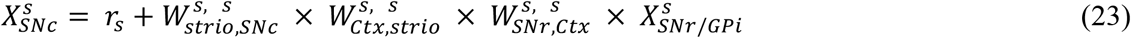

To mitigate this, constants (*C*) are introduced into Equation 22 to stabilize the SNr/GPi output until the synaptic weights are modified by plasticity, as shown in Equation 24.

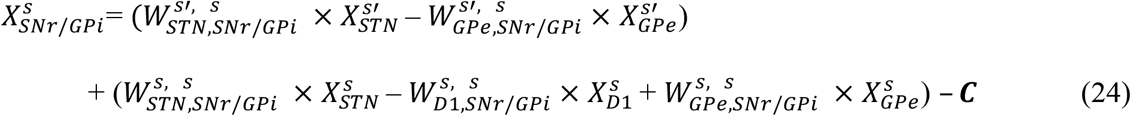

### Transition of inter-module and intra-module projection

To illustrate the transitions of the intra- and inter-module components, we used the activities in the BG-module corresponding to the state two steps ahead of the goal. The inputs from the intra- and inter-module projections were calculated using the following formula:

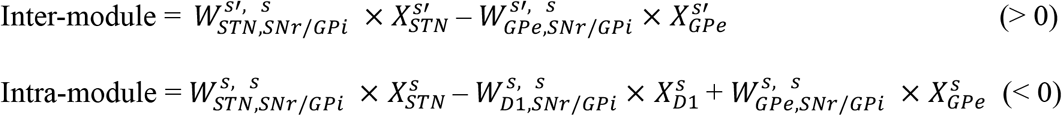

Because the input from the intra-module component was summed to a negative value, the inverted values were calculated, as shown in Figs. 3G and 3H.

### The 2D-maze task

The maze consists of 81 states arranged in a 9 × 9 grid. Excluding the 41 impassable states, the agent explored a path from the start (S) to the goal (G) within the remaining 40 states (Fig. 2A).

The agent chose one of four possible directions (up, down, left, or right) to move in each step. Upon reaching the goal, the agent receives a reward of 1 and returns to the start. The agent starts from the top-left state, and the goal is always in the bottom-right state.

### Conflict task

A 5 × 5 maze was used in the conflict task. Two types of rewards were set for the task: high cost-high return and low cost-low return. The high-cost, high-return reward was placed at a position requiring a minimum of six steps from the start and offered a reward value of +3. The low-cost, low-return reward was placed at a position requiring a minimum of five steps from the start and offered a reward value of +1. Reaching either reward finished the episode, and the agent was reset to the start position. Again, as this task was performed using a state-value-based model, we applied an arbitrary softmax policy for action selection.

During the conflict task, we changed the strengths of the inter- and intra-module projections (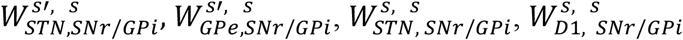, and 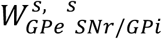.) to make γ^(inter/intra)^ range between 0.1 and 0.9. For each combination of parameters, we performed 300 episodes of learning across 200 sessions. At the end of each session, we calculated the selection probabilities of the two types of reward based on the last 100 episodes. This process of learning and sampling reward selection probabilities was repeated for 200 sessions, and the mean and variance of the number of times each reward was obtained were recorded. The number of times each reward was obtained was then divided by the total number of sessions, converted to a selection probability, and illustrated. The strength of each projection weight was set as listed in Table 4. Weights were set to yield γ^(inter/intra)^ values ranging from 0.1 to 0.9 in increments of 0.1.

### TD-learning model

In state-value-based RL (V-learning), the expected cumulative reward for a given state V(*s*_*t*_) is iteratively updated to approximate an optimal policy. The updates to V(*s*_*t*_) are based on the following equation:

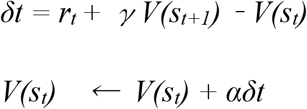

Firstly, the TD-error was calculated, then the state value was updated using the TD-error, where *r*_*t*_ is the reward obtained at time *t, γ* is the discount factor (0 < *γ* < 1), and *α* is the learning rate (*α* ∈ [0, 1]). For further details on the derivation of these equations, see Sutton and Barto (2018)^10^. We used the same action selection policy as the tiled network, that is, the epsilon-greedy policy (epsilon = 0.1).

For the conflict task, we applied various discount rates γ ranging from 0.1 to 0.9 in increments of 0.1. Each agent performed 200 sessions of learning identical to the tiled network, and we counted the number of times each reward was obtained to calculate the selection probabilities. We used the softmax function for the action-selection policy, which was consistent with the policy used in the tiled network.

For action value-based RL (Q-learning), the expected cumulative reward for a given state-action pair Q(*s*_*t*_, *a*_*t*_) is iteratively updated to approximate the optimal policy. The updates to Q(*s*_*t*_, *a*_*t*_) are based on the following equation:

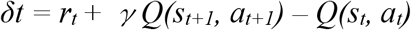

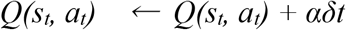

where γ is the discount factor (0 < γ < 1), and α is the learning rate (α ∈ [0, 1]). This method, known as Q-learning with a TD error, provides a way to approximate the correct Q-values by incrementally updating them based on the TD error. For further details on the derivation of these equations, see Sutton and Barto (2018)^10^. The specific parameters used to implement these models are listed in Table 3.

**Table 3.** V-learning and Q-learning parameters.

| Constant | Value | Description |
| --- | --- | --- |
| $\alpha$ | 0.05 | Learning rate |
| $\epsilon$ | $0.5 * (1 / (ep + 1e-8))$ | Probability of random action in epsilon-greedy method.<br>ep represents the episode count within the session. |
| $\gamma$ | 0.1–0.9 | Discount rate. Normally 0.9 except during conflict task. |

**Table 4.**
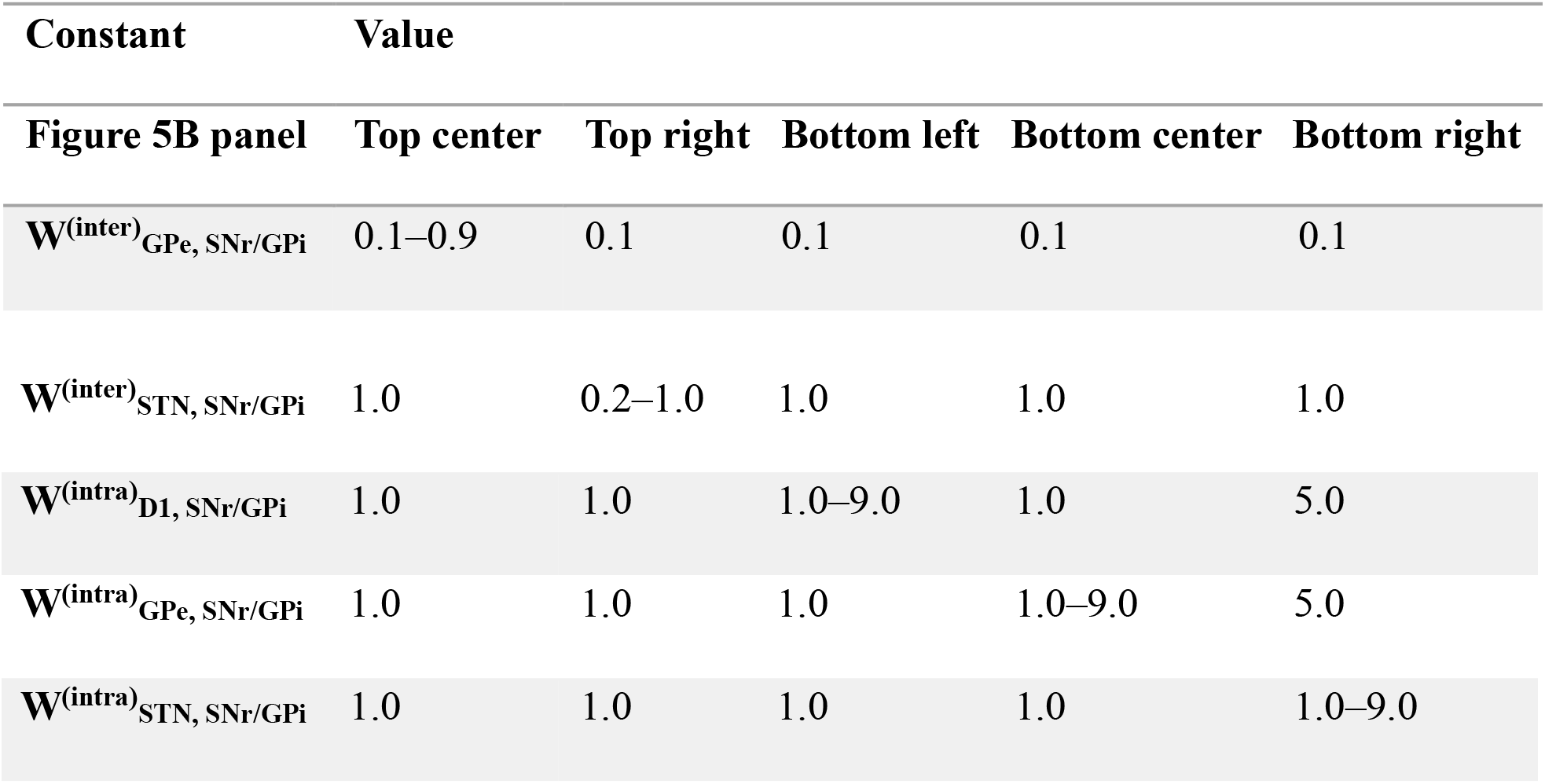
Weight parameters during the conflict task.

### Action value model of horizontal tiled network

In the action value model of the tiled network, we connected inter-module projections between BG-modules corresponding to state-action pairs (*s, a*), the modules corresponding to the next state (*s’*), and all possible actions (*a*_*1*_, *a*_*2*_, *…, a*_*n*_) in that state. Specifically, inter-module projections exist between the BG-module for (*s, a*) and the BG-modules for each (*s’, a*_*1*_), (*s’, a*_*2*_), (*s’, a*_*n*_). In the maze task, the next state (*s’*) for the current state (*s*) is always the same as long as the same action is taken. However, in the cart-pole task, the next state (*s’*) can vary even if the current state (*s*) and action (*a*) pairs are the same. Therefore, in the cart-pole task, action (*a*) was determined first. The inter-module projections were then computed once the next state (*s’*) was uniquely determined. The activity of each nucleus was calculated using the following equation:

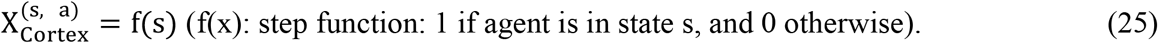

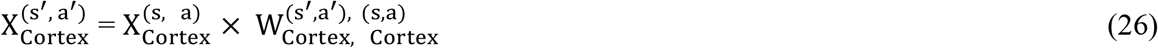

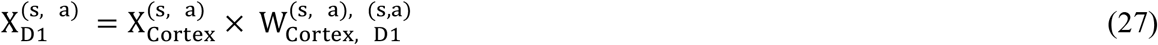

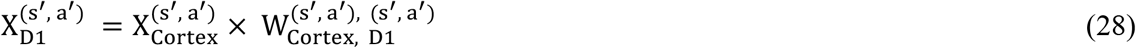

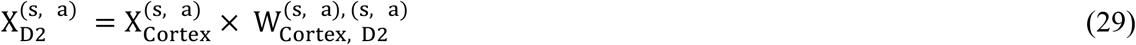

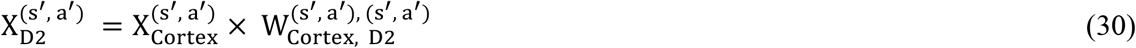

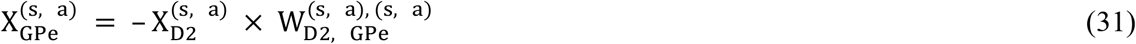

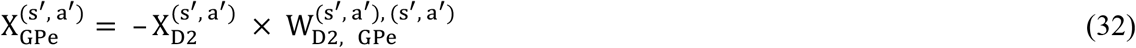

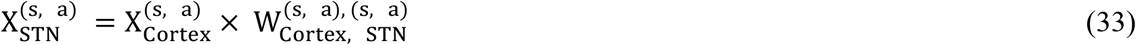

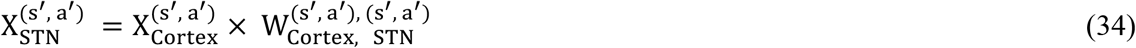

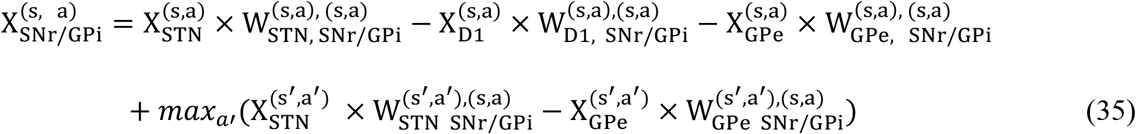

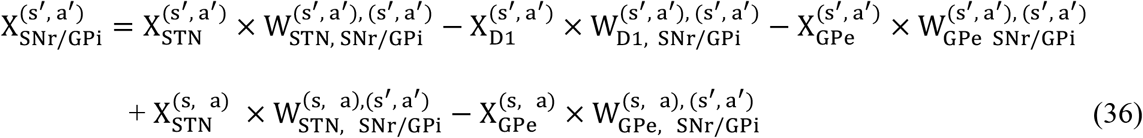

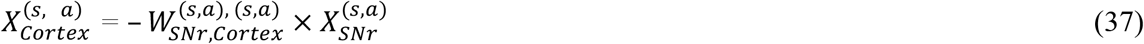

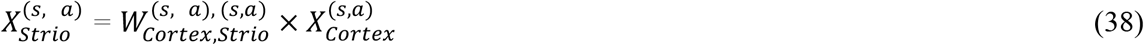

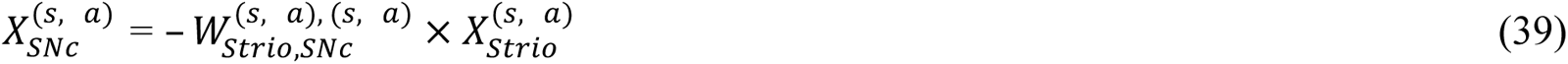

In the context of the current state, the equations include the activity and weights within the BG-modules corresponding to all possible state-action pairs (*s, a*). For the next possible state *s’*, the equations include calculations for all possible state-action pairs (*s’, a’*). The action-value model of the tiled network learns the maze by changing the activity and weight of each nucleus. The learning process followed the algorithm outlined below:

1. Input to the cortex: Input 1 to the cortical neurons in the BG-module corresponding to the current state-action pair 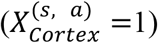.
2. Feedforward Computation: Perform feedforward computations for the BG-module at the current state. Update the activities of the neural nuclei.
3. Action selection: The activities of the STN in the BG-modules corresponding to the current state are compared. The action is selected based on the BG-module with the highest STN activity (highest value) 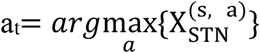. When an agent selects an action, the next state can be determined.
4. State Transition: Transition to the new state (*s*_*t*_ ← *s*_*t+1*_). Execute steps 1 and 2 in the next state.
5. Weight update: Update the weights based on the activity of each nucleus.
6. Reset Activity: Reset the activity levels in all BG-modules.
7. Repeat steps 1 to 6 until the agent reaches the goal (1 episode).

We defined the process from the start to reaching the goal as one episode, and 200 episodes as one session. At the end of each session, all the weights in the BG-modules were reset to their initial values for relearning.

### Cart-pole task

The cart-pole task was implemented using the Gym library provided by OpenAI. Gym offer various environments, and actions can be input into the environment to obtain observations, including the next state and reward. For the cart-pole task, default values for the environmental variables were used. After taking one step, a reward of +1 was given if the pole did not fall, and a reward of -200 was given if the pole fell. The variables in this task were continuous; therefore, we discretized each continuous variable into six bins, resulting in 6^4^ = 1296 possible states.

The action value model of the tiled network learns the cart-pole task by varying the activity and weight of each nucleus. The learning process followed the algorithm outlined below:

1. Input to Cortex: Input 1 to the cortex in the BG-module corresponding to the current state-action pair 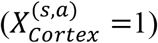.
2. Feedforward Computation: Perform feedforward computations for the BG-module corresponding to the current state. Update the activities of the neural nuclei.
3. Action selection: The activities of the STN in the BG-modules corresponding to the current state are compared. Select the action with the highest activity (highest value) 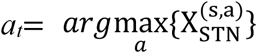, and determine the next state transition.
4. State Transition: Transition to the new state (*s*_*t*_ ← *s*_*t+1*_). Execute steps 1 and 2.
5. Feedforward Computation: Perform feedforward computations for the BG-module corresponding to the previous state. Update the weights of the neural nuclei.
6. Reset Activity: Reset the activity levels in all BG-modules.
7. Repeat steps 1 to 7 until the agent reaches the goal (1 episode).

Using the above procedure, an episode was defined as either successfully completing 200 actions without the pole falling or failing to complete it. Each session consisted of 800 episodes. At the end of each session, all synaptic weights in the basal-ganglia units were reset to their initial values, and the agent underwent relearning.

In this task, the tiled network determines the actions for each step, which are then input into the gymnasium environment to obtain the next observation. This observation is subsequently input back into the tiled network to determine the next action, continuing the learning process. When the pole fell, the episode immediately ended, and a negative reward of -200 was delivered. Conversely, if the agent could keep the pole balanced for 195 steps, the episode ended, and a positive reward of +200 was given. We performed 200 sessions of learning for both the tiled network and Q-learning model, with each session consisting of 800 episodes. The performance of the two models was compared.

### Heatmap for Q-value visualization

In the cart-pole task, the entire state-action space can be represented as (*s*_1_, *s*_2_, *s*_3_, *s*_4_, a) ∈ (***S*** × ***A***), with a BG-module tiled over each element in the state-action space. where *s*_1_ represents the cart position, *s*_2_ the pole angle, *s*_3_ the cart velocity, *s*_4_ the pole tip velocity, and a the action (right or left). For each state (*s*_1_, *s*_2_, *s*_3_, *s*_4_), two BG-modules correspond to the two actions (left or right). We visualized the highest STN activity among the two BG-modules:

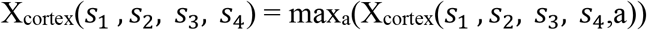

To visualize the STN activities in a two-dimensional heatmap, we averaged the STN activities across dimensions *s*_3_ (cart velocity) and *s*_4_ (pole tip velocity), focusing on the two variables *s*_1_ (cart position) and *s*_2_ (pole angle):

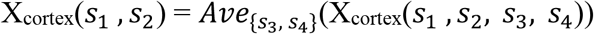

The resulting X_cortex_(*s*_1_, *s*_2_) was visualized as a heat map. For comparison, we trained a TD-error-based Q-learning model. We also performed the same dimensionality reduction process as the Q-values and visualized it as a two-dimensional heatmap.

## Data availability

No datasets were generated or analyzed during the current study.

## Computer code

The custom codes used in this study have been deposited in the database (Zenodo) and are publicly accessible at doi: 10.5281/zenodo.15581467.

## Acknowledgments

We thank Dr. Daniel J. Gibson for critically reading the manuscript and providing excellent suggestions for analysis. This research was supported by JST SPRING (JPMJSP2138 to K.H.) and KAKENHI (18H04945 and 22K18662 to T.K.).

## Author contributions

[Conceptualization] K.H. and T.K. designed the study. [Methodology] K.H., R.T., T.K., S.O., and T.K. developed the modeling framework, including the selection of algorithms and computational techniques. [Formal Analysis] K.H., R.T., T.K., and S.O. developed the actual code. [Visualization] R.T., T.K., and S.O. created figures, graphs, and other visual representations. [Manuscript Writing] K.H. wrote the original draft. K.H., R.T., T.K., S.O., and T.K. edited and reviewed the manuscript. [Supervision and Funding] T.K. supervised the project and secured funding. All authors have reviewed and approved the final manuscript.

## References

1. Samejima K, Ueda Y, Doya K, Kimura M. Representation of action-specific reward values in the striatum. Science. 2005; 310: 1337–1340.

2. Schultz W, Dayan P, Montague PR. A neural substrate of prediction and reward. Science. 1997; 275: 1593–1599.

3. Cohen JY, Haesler S, Vong L, Lowell BB, Uchida N. Neuron-type-specific signals for reward and punishment in the ventral tegmental area. Nature. 2012; 482: 85–88.

4. Flaherty AW, Graybiel AM. Two input systems for body representations in the primate striatal matrix: experimental evidence in the squirrel monkey. J Neurosci. 1993; 13: 1120–1137.

5. Nambu A, Tokuno H, Takada M. Functional significance of the cortico-subthalamo-pallidal ‘hyperdirect’ pathway. Neurosci Res. 2002; 43: 111–117.

6. Mink JW. The basal ganglia: focused selection and inhibition of competing motor programs. Prog Neurobiol. 1996; 50: 381–425.

7. Nambu A, Tokuno H, Hamada I, Kita H, Imanishi M, Akazawa T, Ikeuchi Y, Hasegawa N. Excitatory cortical inputs to pallidal neurons via the subthalamic nucleus in the monkey. J Neurophysiol. 2000; 84: 289–300.

8. Nambu A. Somatotopic organization of the primate Basal Ganglia. Front Neuroanat. 2011; 5: 26.

9. Ozaki M, Sano H, Sato S, Ogura M, Mushiake H, Chiken S et al. Optogenetic activation of the sensorimotor cortex reveals ‘local inhibitory and global excitatory’ inputs to the basal ganglia. Cereb Cortex N.Y.N 1991. 2017; 27: 5716–5726.

10. Sutton RS, Barto AG. Reinforcement Learning: an Introduction. xxii. 2nd edn. (The MIT Press, Cambridge, MA, 2018), p 526.

11. Grillner S, Robertson B. The basal ganglia downstream control of brainstem motor centres-- an evolutionarily conserved strategy. Curr Opin Neurobiol. 2015; 33: 47–52.

12. Alexander GE, DeLong MR, Strick PL. Parallel organization of functionally segregated circuits linking basal ganglia and cortex. Annu Rev Neurosci. 1986; 9: 357–381.

13. Albin RL, Young AB, Penney JB. The functional anatomy of basal ganglia disorders. Trends Neurosci. 1989; 12: 366–375.

14. Hernández-López S, Bargas J, Surmeier DJ, Reyes A, Galarraga E. D1 receptor activation enhances evoked discharge in neostriatal medium spiny neurons by modulating an L-type Ca2+ conductance. J Neurosci. 1997; 17: 3334–3342.

15. Gerfen CR, Surmeier DJ. Modulation of striatal projection systems by dopamine. Annu Rev Neurosci. 2011; 34: 441–466.

16. Surmeier DJ, Ding J, Day M, Wang Z, Shen W. D1 and D2 dopamine-receptor modulation of striatal glutamatergic signaling in striatal medium spiny neurons. Trends Neurosci. 2007; 30: 228–235.

17. Yagishita S, Hayashi-Takagi A, Ellis-Davies GCR, Urakubo H, Ishii S, Kasai H. A critical time window for dopamine actions on the structural plasticity of dendritic spines. Science. 2014; 345: 1616–1620.

18. Graybiel AM, Hickey TL. Chemospecificity of ontogenetic units in the striatum: demonstration by combining [3H]thymidine neuronography and histochemical staining. Proc Natl Acad Sci U S A. 1982; 79: 198–202.

19. Gerfen CR. The neostriatal mosaic: compartmentalization of corticostriatal input and striatonigral output systems. Nature. 1984; 311: 461–464.

20. Gerfen CR. The neostriatal mosaic. I. Compartmental organization of projections from the striatum to the substantia nigra in the rat. J Comp Neurol. 1985; 236: 454–476.

21. Jimenez-Castellanos J, Graybiel AM. Subdivisions of the primate substantia nigra pars compacta detected by acetylcholinesterase histochemisty. Brain Res. 1987; 437: 349–354.

22. Zhang J-C, Lau P-M, Bi G-Q. Gain in sensitivity and loss in temporal contrast of STDP by dopaminergic modulation at hippocampal synapses. Proc Natl Acad Sci U S A. 2009; 106: 13028–13033.

23. Frémaux N, Gerstner W. Neuromodulated spike-timing-dependent plasticity, and theory of three-factor learning rules. Front Neural Circuits. 2015; 9: 85.

24. Schultz W, Apicella P, Ljungberg T. Responses of monkey dopamine neurons to reward and conditioned stimuli during successive steps of learning a delayed response task. J Neurosci. 1993; 13: 900–913.

25. Krausz TA, Comrie AE, Kahn AE, Frank LM, Daw ND, Berke JD. Dual credit assignment processes underlie dopamine signals in a complex spatial environment. Neuron. 2023; 111: 3465–3478.e7.

26. Kim MJ, Gibson DJ, Hu D, Yoshida T, Hueske E, Matsushima A, Mahar A, Schofield CJ, Sompolpong P, Tran KT, Tian L, Graybiel AM. Dopamine release plateau and outcome signals in dorsal striatum contrast with classic reinforcement learning formulations. Nat Commun. 2024; 15: 8856.

27. Flaherty AW, Graybiel AM. Input-output organization of the sensorimotor striatum in the squirrel monkey. J Neurosci. 1994; 14: 599–610.

28. DeLong M, Wichmann T. Changing views of basal ganglia circuits and circuit disorders. Clin EEG Neurosci. 2010; 41: 61–67.

29. Guthrie M, Leblois A, Garenne A, Boraud T. Interaction between cognitive and motor cortico-basal ganglia loops during decision making: a computational study. J Neurophysiol. 2013; 109: 3025–3040.

30. Foster NN, Barry J, Korobkova L, Garcia L, Gao L, Becerra M et al. The mouse cortico-basal ganglia-thalamic network. Nature. 2021; 598: 188–194.

31. Hebb DO. The Organization of Behavior; a Neuropsychological Theory. xix. (John Wiley & Sons, Oxford, England, 1949), p 335.

32. Reynolds JN, Hyland BI, Wickens JR. A cellular mechanism of reward-related learning. Nature. 2001; 413: 67–70.

33. Kreitzer AC, Malenka RC. Endocannabinoid-mediated rescue of striatal LTD and motor deficits in Parkinson’s disease models. Nature. 2007; 445: 643–647.

34. Graybiel AM, Ragsdale CW, Yoneoka ES, Elde RP. An immunohistochemical study of enkephalins and other neuropeptides in the striatum of the cat with evidence that the opiate peptides are arranged to form mosaic patterns in register with the striosomal compartments visible by acetylcholinesterase staining. Neuroscience. 1981; 6: 377–397.

35. Doya K. Metalearning and neuromodulation. Neural Netw. 2002; 15: 495–506.

36. Miyazaki KW, Miyazaki K, Tanaka KF, Yamanaka A, Takahashi A, Tabuchi S et al. Optogenetic activation of dorsal raphe serotonin neurons enhances patience for future rewards. Curr Biol. 2014; 24: 2033–2040.

37. Schweighofer N, Bertin M, Shishida K, Okamoto Y, Tanaka SC, Yamawaki S et al. Low-serotonin levels increase delayed reward discounting in humans. J Neurosci. 2008; 28: 4528– 4532.

38. Lazaridis I, Crittenden JR, Ahn G, Hirokane K, Wickersham IR, Yoshida T et al. Striosomes control dopamine via dual pathways paralleling canonical basal ganglia circuits. Curr Biol. 2024; 34: 5263–5283.e8.

