## Supplemental Text for "Horizontally Tiled Network of Cortico-Basal Ganglia Modules Performs Reinforcement Learning"

### Supplementary information

**Appendix 1. Theoretical confirmation of intra- and inter-module projection encode Vs and Vs' individually.**

It is possible to theoretically confirm that the two pathways projecting to the SNr/GPi could encode information equivalent to *Vs* and *Vs'*. When the agent reached the goal and received a reward, the dopamine activity ($X^{s}_{SNc}$) in the goal state increased. Subsequently, dopamine-dependent Hebbian plasticity led to an increase in the Cortex-STN and Cortex-striatal D1 weights, while the Cortex-striatal D2 weight decreased (Equations 17-19). Therefore, the cortex and STN were positively correlated with the state value, whereas the D2 was negatively correlated. As the striatal D2-GPe is an inhibitory projection, GPe has a positive correlation with the state value. Thus, the relationship between the activity of each nucleus and the state value can be expressed as follows:

$$X^{s}_{STN} = \alpha_{STN} V^{s}$$

$$X^{s}_{D1} = \alpha_{D1} V^{s}$$

$$X^{s}_{GPe} = \alpha_{D2} V^{s}$$

Here, $\alpha_{STN}$, $\alpha_{D1}$, $\alpha_{D2}$ were constant parameters, and in this study, all the parameters were set to 1.0, so these parameters were omitted in the subsequent equations. Therefore, the input to the SNr/GPi can be expressed as follows:

$$X^{s}_{SNr} = \text{(inter-module projection)} + \text{(intra-module projection)}$$

$$= (W^{s', s}_{STN,SNr/GPi} \times X^{s'}_{STN} - W^{s', s}_{GPe,SNr/GPi} \times X^{s'}_{Gpe})$$

$$+ (W^{s,\ s}_{STN,SNr} \times X^{s}_{STN} - W^{s,(intra)}_{D1,SNr} \times X^{s}_{D1} + W^{s,(intra)}_{GPe,SNr} \times X^{s}_{Gpe})$$

$$= (W^{s\prime,(inter)}_{STN,SNr} \times V(s') - W^{s\prime,(inter)}_{GPe,SNr} \times V(s'))$$

$$+ (W^{s,(intra)}_{STN,SNr} \times V(s) - W^{s,(intra)}_{D1,SNr} V(s) - W^{s,(intra)}_{GPe,Snr} \times V(s))$$

$$= (W^{s\prime,(inter)}_{STN,SNr} - W^{s\prime,(inter)}_{GPe,Snr})V(s')$$

$$- (-W^{s,\ s}_{STN,\ SNr/GPi} + W^{s,\ s}_{D1,\ SNr/GPi} + W^{s,\ s}_{GPe\ SNr/GPi})V(s)$$

From this, the SNc activity could be written as following:

$$SNc = r_{(s)} + (W^{s',(inter)}_{STN,SNr} - W^{s',(inter)}_{GPe,SNr})V(s')$$

$$- (-W^{s,\ s}_{STN,\ SNr/GPi} + W^{s,\ s}_{D1,\ SNr/GPi} + W^{s,\ s}_{GPe\ SNr/GPi})V(s)$$

$$\propto\ r_{(s)} + \gamma^{(inter/intra)} V(s') - V(s)$$

$$=\ \delta t$$

In the above equation, $\gamma^{(inter/intra)}$ could be considered as the discount factor. Under the constraint of $0 < \gamma^{(inter/intra)} < 1$ and $(W^{s,(intra)}_{STN,SNr} - W^{s,(intra)}_{D1,SNr} - W^{s,(intra)}_{GPe.Snr}) = -1$, making the SNc activity equivalent to the TD-error.
